# Single-cell transcriptomic profiling of the whole colony of *Botrylloides diegensis*: Insights into tissue specialization and blastogenesis

**DOI:** 10.1101/2024.07.21.604516

**Authors:** Berivan Temiz, Michael Meier, Megan J. Wilson

## Abstract

*Botrylloides diegensis* is a colonial ascidian that has been the focus of developmental, evolutionary, and regeneration research. In this study, we performed single-cell RNA sequencing (scRNA-seq) of an entire *B. diegensis* colony, including zooids, buds, and vascular tunics, to resolve cellular heterogeneity and identify cell and tissue markers. We identified 29 major cell clusters within the colony and used *in situ* hybridization to examine the spatial expression of cluster marker genes. Numerous tissue types were identified at the molecular level, including blood cells and zooid tissues such as the branchial epithelium, stomach, and endostyle. Distinct cluster markers were identified for specific regions of the stomach epithelium, highlighting the specialization of these regions and the strength of using scRNA-seq to explore their functionality. Cell trajectory projections highlighted the early appearance of progenitor clusters, whereas more differentiated zooid-related tissues appeared later in the developmental path. This study provides a valuable resource for understanding the development, tissue function, and regeneration of *B. diegensis.* This demonstrates the power of scRNA-seq to define cell types and tissues in complex colonial organisms.

**Summary statement:** Single-cell RNA sequencing of *Botrylloides diegensis* revealed cellular heterogeneity, identified 29 major cell clusters, and provided insights into tissue specialization and blastogenesis.

## Introduction

Single-cell RNA sequencing (scRNA-seq) can quantify the total transcriptome of individual cells in the tissue/organism of interest. This information can be used to classify, characterize, and distinguish each cell at the mRNA level, allowing for the identification of different cell types, developmental states, and trajectories. Many studies have mapped the transcriptomic composition of diverse animals, including invertebrates such as planaria, sponges, *Hydra*, and ascidian species. These studies are essential to understand how cell types differ from each other through the activation or repression of specific pathways; therefore, they can explain how cells possess pluripotency, commit to becoming specific biological units, access categorical morphology, and become a part of tissue during development or regeneration.

Colonial ascidians are sessile marine chordates (Fig. 1A), which can be categorized into three main parts:1) zooids, buds, and budlets as the asexually developing body; 2) the vascular network, including veins, blood cells, and vascular termini/ampullae; and 3) tunic, the gelatinous matrix covering all colonies (Berrill, 1947a; Blanchoud et al., 2017) (Fig. 1B and C). At least 11 blood cell types have been identified in the hemolymph and are grouped into five cell types:1) undifferentiated cells, hemoblasts, and differentiating cells; 2) immunocytes, hyaline amebocytes, macrophage-like cells, granular amebocytes, and morula cells; 3) transport cells, compartment amebocytes, and compartment cells; 4) mast cell-like cells or granular cells; and 5) storage cells, pigment cells, or nephrocytes(Blanchoud et al., 2017; Cima et al., 2001; Hirose et al., 2003; Wright and Ermak, 1982). The zooid, as a filter-feeding individual, consists of various anatomical structures, including the atrial epithelium, atrial siphon, branchial epithelium, bud, endostyle, gonads, intestine, neural complex, peribranchial sacs, pericardium, pyloric gland, stigmata, and the stomach (Fig. 1D and E) (Anselmi et al., 2022; Berrill, 1947a; Blanchoud et al., 2017; Holland, 2016; Kawamura and Sunanaga, 2010). During asexual reproduction and blastogenesis, a new zooid is produced by budding from the atrial epithelium of the parental zooid (Berrill, 1947a; Manni and Burighel, 2006; Mukai et al., 1978). This budding process, also known as blastogenesis or palleal budding, is morphologically similar to whole-body regeneration (WBR) (Berrill, 1951; Rinkevich et al., 2007). Thus, studying intact whole colonies will help capture the critical aspects of their development, tissue function, and regeneration.

**Figure 1.**
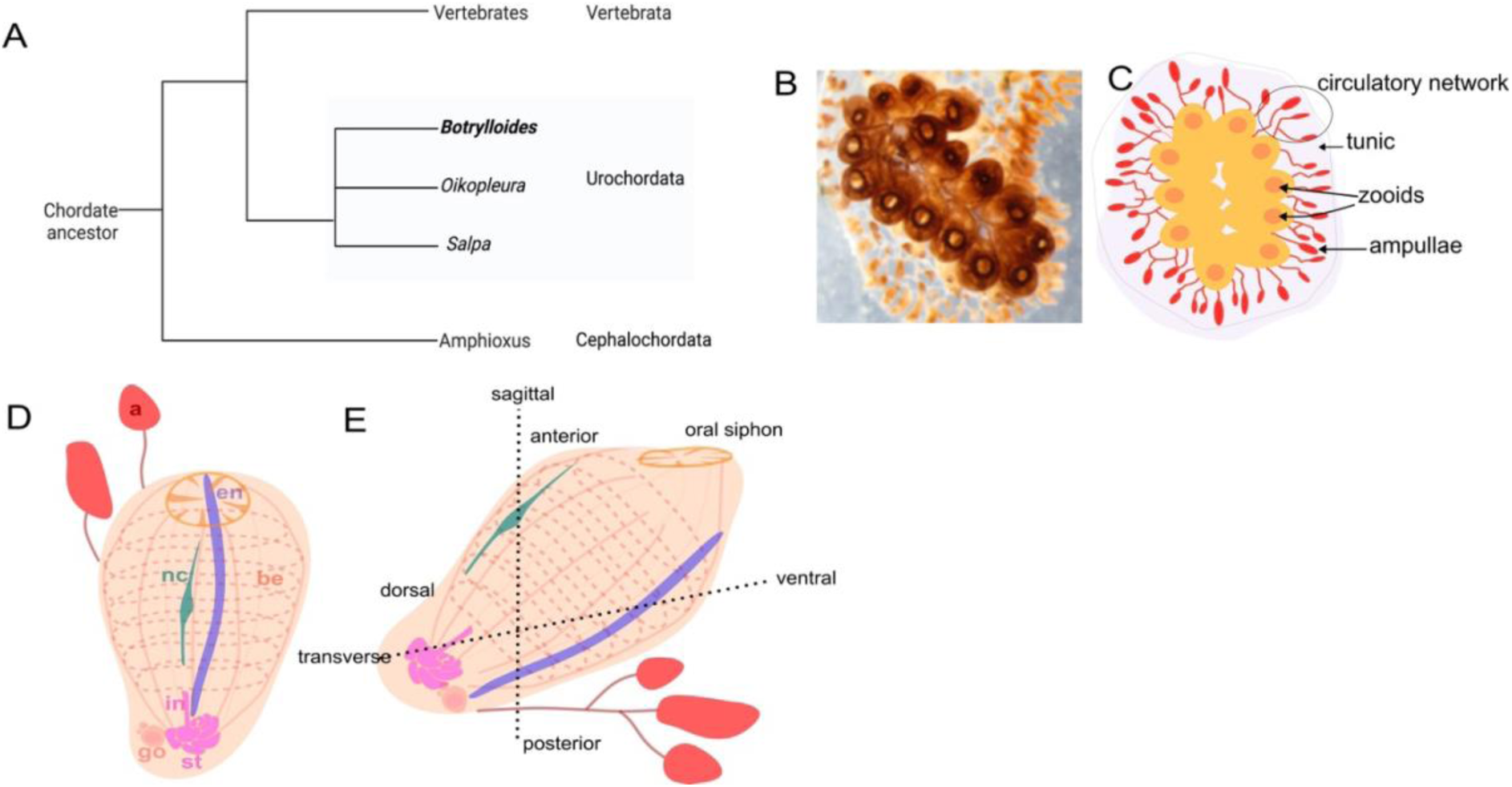
The study system: *Botryllodies diegensis.* A. Simplified phylogeny represents the position of tunicates as the closest chordate group to vertebrates. B. Example *B. diegensis* colony attached to a glass slide. C. Orange morphs of *the B. diegensis* colony are illustrated. The red lines indicate shared circulatory networks. The vascular termini (ampulla) are elliptical vascular ends. The individuals (zooids) are lined side by side in a ladder-like arrangement in the gelatinous matrix (tunic). D and E. Simplified zooid anatomy, dorsal (D) and lateral (E) views. Abbreviations: intestine (in), stomach (st), gonad (go), neural complex (nc), endostyle (en), branchial epithelium (be).

Several sc-RNA-seq studies have been performed with the solidary ascidian *Ciona intestinalis*, mainly during the embryonic stage, showing the conservation of developmental and functional programs in chordate evolution (Cao et al., 2019; Horie et al., 2018; Sharma et al., 2019; Zhang et al., 2020). *B. diegensis* is an emerging model for development, regeneration, and stem cell studies owing to its small genome size, short replenishment period, and excellent example of WBR (Blanchoud et al., 2018). Therefore, investigating the composition of mature colonies is crucial to understand the cell types and tissues of *B. diegensis*.

This study aimed to develop a protocol and resource for *B. diegensis* to identify cell and tissue markers. Single-cell transcriptomic profiling of a mature (blastogenic stage A) *B. diegensis* colony was performed. Consequently, several cell and tissue types were separated based on their unique gene expression profiles, and their spatial expression was validated using *in situ* hybridization.

## Results and Discussion

*B. diegensis* colonies were obtained from a local marina and kept in the laboratory. We performed scRNA-seq using acetic acid-methanol (ACME) tissue dissociation and fixation methods (Fig. 2A). Cells from a *B. diegensis* colony at stage A of the blastogenic cycle were fixed and dissociated using ACME (Fig S1 A-B). This was followed by FACS sorting to remove cell aggregates and debris (Fig.S1E-L). Chromium^TM^ Single-cell 3’ (10X Genomics) was used for single-cell barcoding. The barcoded reads were sequenced in both directions for paired-end sequencing. Library quality control was performed using FastQC software (Fig. S2). High-quality reads were mapped back to the reference genome using STARSolo software. The summary statistics for mapping are provided in Table S1. A single-cell transcriptome library containing approximately 58 million reads was generated. The percentage of the aligned reads with valid barcodes was 96%. The rate of reads that mapped to unique genes was 73%. In total, 6353 cells were detected, with a mean of 1586 UMIs and 481 genes per cell. A total of 13621 genes were identified (Table S1).

**Figure 2.**
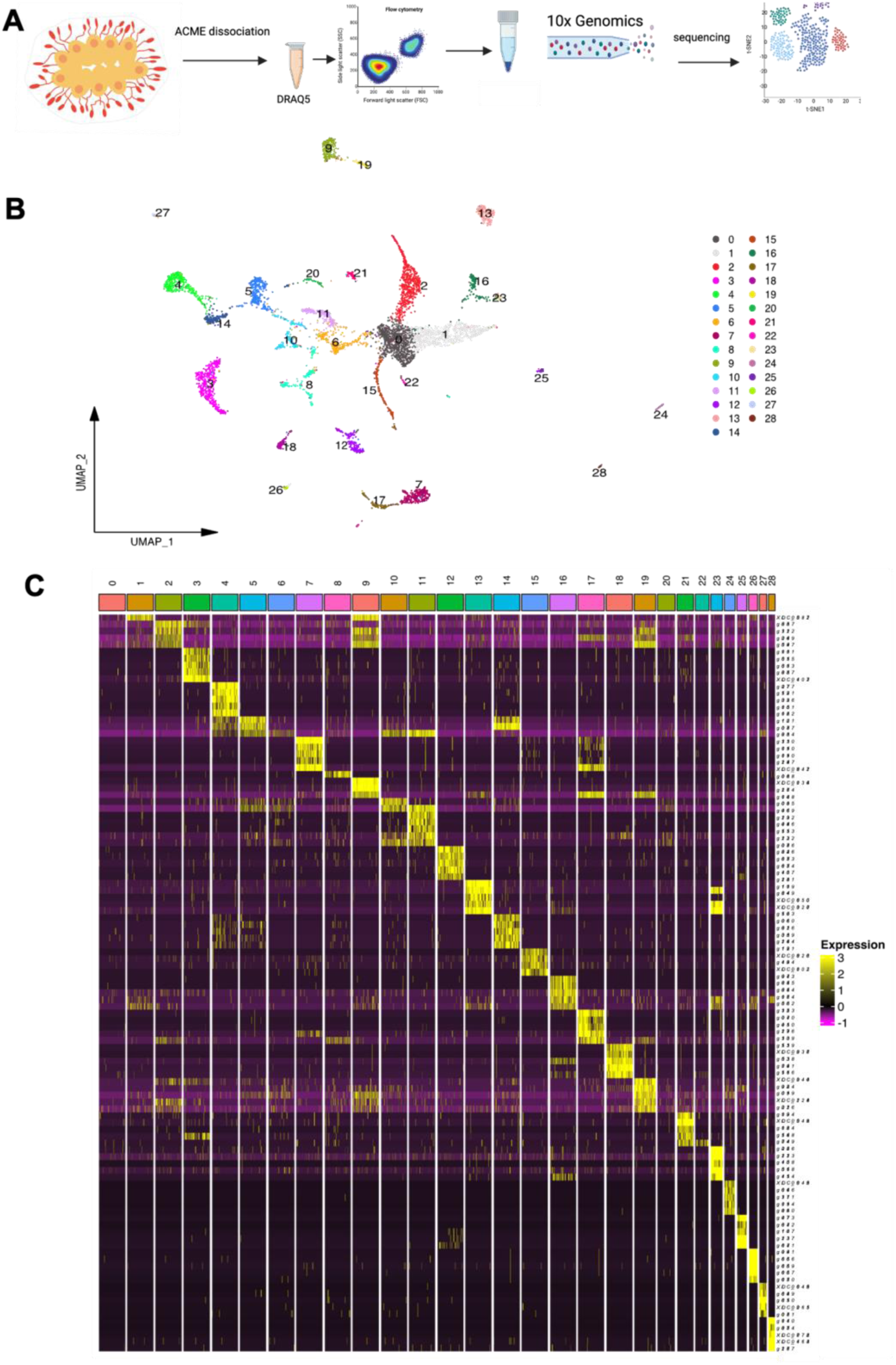
Overview of the experimental pipeline and clustering results. A. Single cells were prepared using the ACME maceration method (Garcia-Castro et al., 2021). Cells were stained with DRAQ5 and DRAQ5+ cells were sorted by FACS. Cells were captured in droplets using a 10X chromium system. Single-cell libraries were prepared and sequenced. After mapping the transcripts to the genome, clustering analysis was performed to identify the cell types. B. UMAP clustering analysis revealed 29 single-cell clusters. Each cluster was color-coded. C. Heatmap of the top five marker genes in each cluster. Abbreviations: acetic-methanol (ACME).

Cluster analysis was performed to group single-cell transcriptomes based on similarity using the Seurat software. Genes showing high variation between cells were calculated to obtain the best signal in cluster differences, and 2,000 genes were included by default. Twenty-nine distinct clusters were identified using nonlinear dimensional reduction with Uniform Manifold Approximation and Projection (UMAP) (Fig 2B). The cell numbers for each cluster are shown in Fig. S3. The highest cell number was observed for cluster 0, with 779 cells, and the lowest cell number was 25 for cluster 28. The mean cell number/cluster ratio was 208. The average number of genes in each cluster was 291.

### Identification of Cluster Marker Genes

The FindMarkers function was used for differential expression analysis within the Seurat package to identify the top genes that were potential markers for each cluster (Fig. 2C). Differentially expressed genes (DEGs) were recorded as fold changes (LFC). The full DEG list for each cluster is provided in File S1. DEGs were annotated to their closest human orthologs. Genes that did not have vertebrate or human matches were also calculated for each cluster (Table S2). Gene ontology (GO) analysis was conducted for each cluster to aid the functional characterization of the clusters. The closest vertebrate orthologs that matched the DEGs were used to generate gene lists for GO analysis (File S2).

### Using single-cell data to identify regional tissue markers

The expression patterns of the top marker genes of Clusters 4, 5, and 14 were determined by *in situ* hybridization (Fig. 3). *Ctrb1* (*g03753*) was one of the most highly expressed transcripts (∼8 LFC) in Cluster 4, which was exclusively present in this cluster compared to other clusters (Fig 3A). *Ctrb1* encodes a serine protease enzyme linked to the acinar-like exocrine glandular cells involved in digestion (Perillo et al., 2016). A probe was designed to determine the cell-type expression of *Ctrb1,* and a strong staining signal was detected in the stomach of the zooid (Fig 3A). These *Ctrb1+* epithelial cells were a subset of cells located in the outer curling of the stomach folds (Fig. 3A). No staining was observed in other tissues or vascular cells in mature colonies (Fig. 3A).

**Figure 3.**
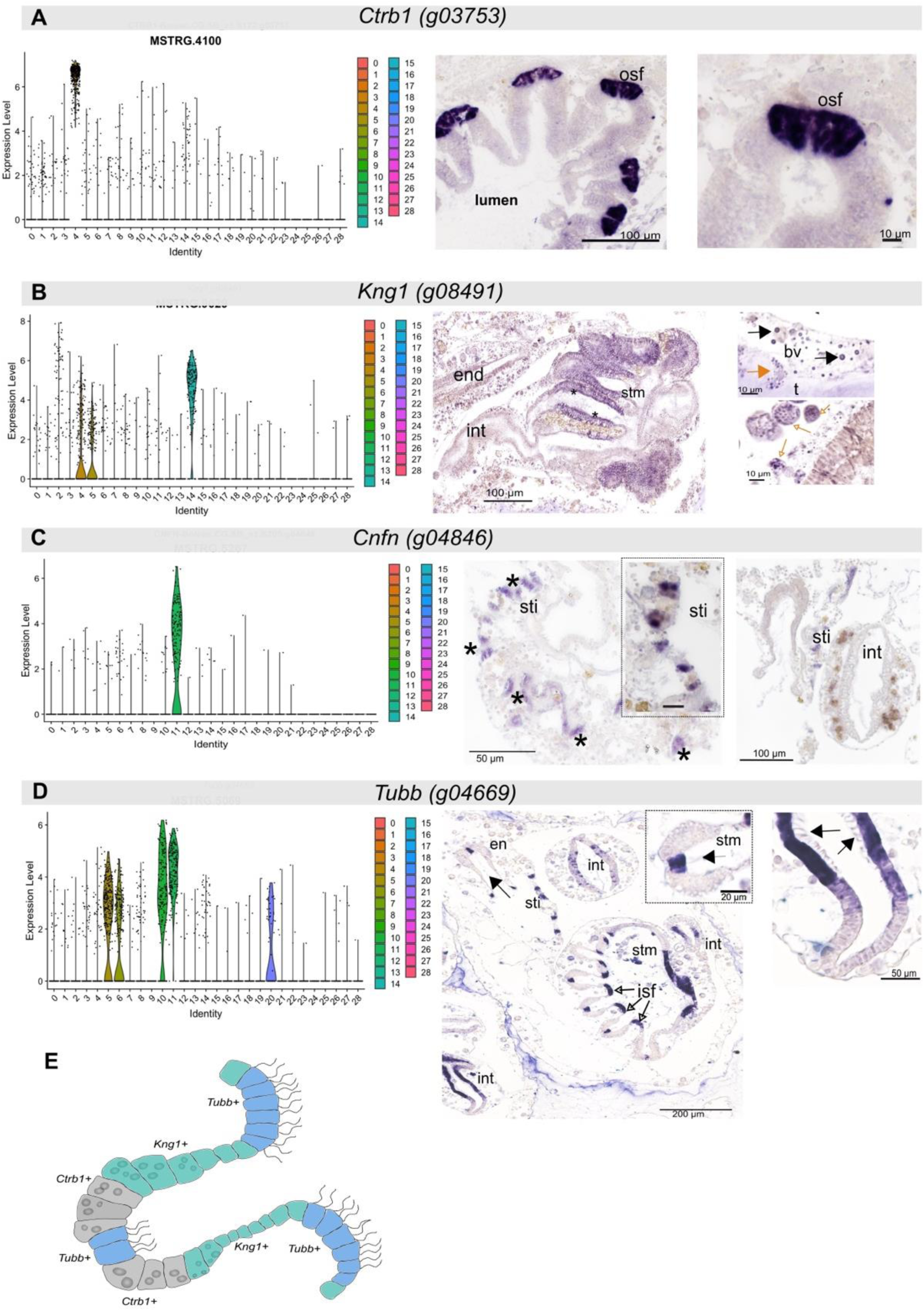
Identification of *B. diegensis* digestive tract clusters. A. Violin plot showing *Ctrb1* as a marker of cluster 4. *In situ* expression of *Ctrb1* mRNA was observed in the cells on the outer side of each stomach fold. B. *Kng1+* cells longitudinal stomach folds. C. Single-cell cluster expression profile for *Cnfn*, g04846, the top marker gene for Cluster 11. The *Cnfn* probe-stained cells are found in rows (asterixis) that line the branchial chamber, known as stigmata cells. D. Cluster expression for *Tubb* was identified as a top marker gene for Clusters 10 and 11, although it was expressed by cells found in several other Clusters*. Tubb* mRNA was found in the stomach epithelium, branchial epithelial cells, and epithelia covering the oocytes/eggs of developing buds. E. Based on the *in situ* results, an Illustration of a stomach fold showing the location of *Tubb+, Ctrb1+* and *Kng1*+ cells in the stomach epithelium. Abbreviations: endostyle (end), stomach (stm), inner stomach fold (isf), outer stomach fold (osf), stigmata (sti), tunic (t), intestine (int).

*Kng1 (g08491)* was identified as a marker for Cluster 14 (File S1) and was present to a lesser extent in Clusters 2, 4, and 5. A 610 bp fragment of *Kng1* was cloned and sequenced for use during *in situ* hybridization. Intense *Kng1* staining was observed within the lateral edges of the stomach via *in situ* hybridization in adult tissue sections (Fig 3B). Thus, Cluster 14 was identified as a part of the stomach epithelium.

Cluster 10 marker g14532, a cornifelin-like gene (*Cnfn)*), showed specific expression in stigmata cells (Fig. 3C). This gene is localized to microtubules in humans, particularly in the epidermis and oral mucosa (Wagner et al., 2019), and is associated with cell-cell adhesion. It contains a cysteine-rich domain known as a PLAC8 domain.

*Tubb* was broadly upregulated in branchial epithelial tissues of the digestive tract (Fig. 3D). Tubb is a tubulin beta protein that functions in the microtubules of the cytoskeleton, controlling cell shape, movement, and transport within the cell (Sewell et al., 2024). Cilia are microtubule-based organelles, and the zone 1 cilium has a different axonemal structure than the other zones (*Ciona* endostyle) (Konno and Inaba, 2020), which may indicate that these cells express distinct combinations of microtubule genes.

GO and pathway analyses were conducted for genes highly expressed in digestive-and branchial-associated tissues (Clusters 4, 5, 10, 11, and 14) (Fig. 4). Clusters 10 and 11 were overrepresented in the pathways linked to cilia assembly and movement (Fig. 4). This aligns with the mRNA expression of *Cnfn* and *Tubb,* markers of cell clusters 11 and 10, respectively, which showed intense staining in cilia-rich cell types (Fig. 3C and D). Annotation of clusters 4, 14, and 5 confirmed the enrichment of biological processes associated with food breakdown, such as metabolic and catabolic oxidoreductase activity and cell secretion (Fig. 4).

**Figure 4.**
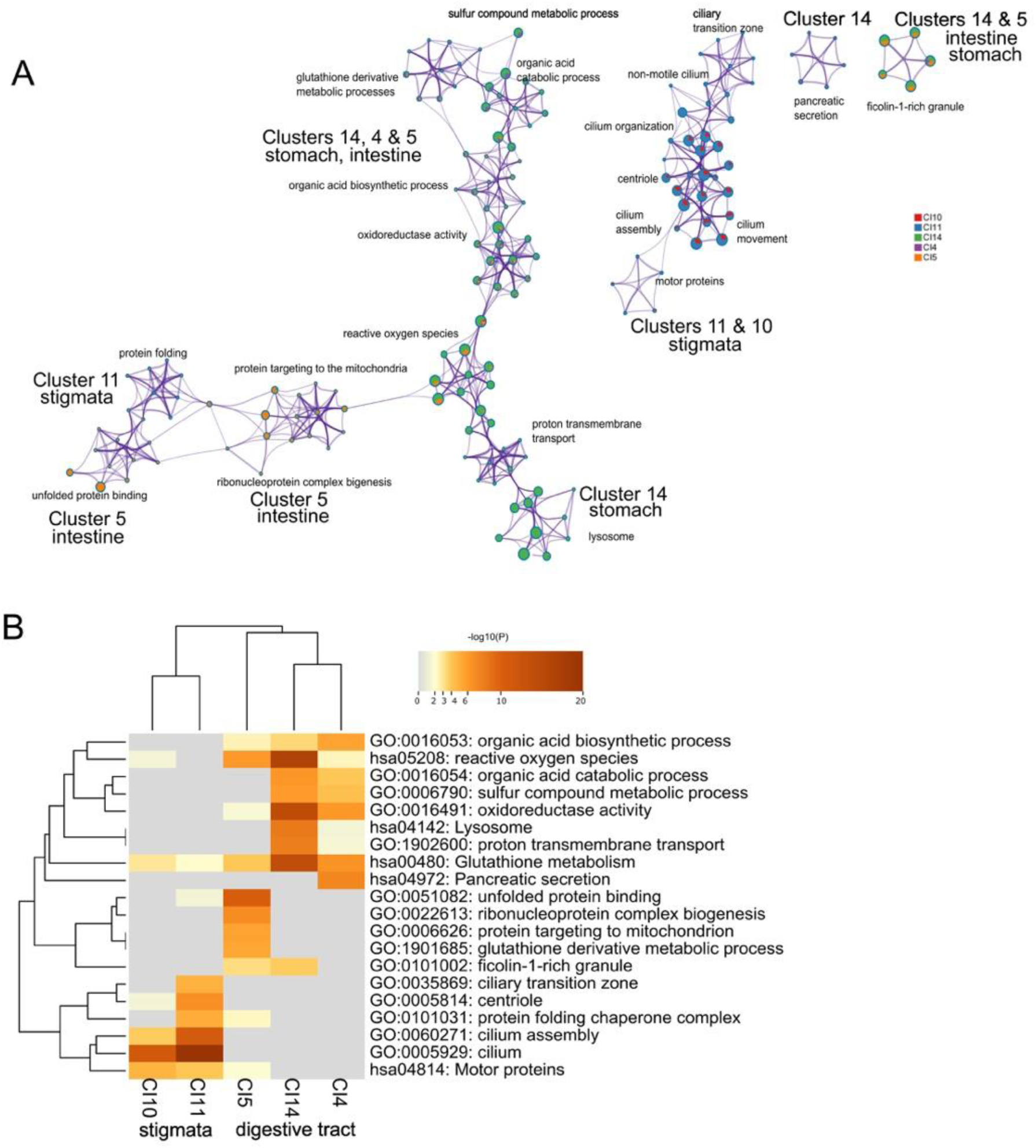
GO and pathway analyses for cell Clusters associated with digestive functions. A. Metascape network of enriched terms, colored by Cluster number and predicted tissue type. Each node is colored according to the number of genes (from each list) associated with that term. B. A clustered heatmap of the top enriched GO terms colored by p-values. Each row represents an enriched GO term, and each column represents the gene list for Clusters 14, 4, 5, 10 and 11.

### Identification of Endostyle clusters

Endostyle, a tissue similar to the pharynx, filters food, synthesizes hormones, and provides immune defences (Takagi et al., 2022). They are also believed to possess stem cell niches or hematopoietic properties (Voskoboynik et al., 2008b). Genes known to be expressed in the endostyle of other ascidian species (*Ciona, Styela*, and *Botryllus*) were selected and identified using the single-cell dataset (Table S4). Clusters 6, 3, and 8 were deemed potential endostyle clusters based on the expression of known endostyle markers. In the ascidian *Ciona*, galectin is expressed in various regions of the endostyle (Parrinello et al., 2015; Parrinello et al., 2017). The *B. diegensis* genome contains multiple *Lgal* genes (File S3) whose transcripts were detected in clusters 3, 4, and 10 (File S3) (Fig. 5A). In *Styela*, *Itnl1/Fcn1* mRNAs were enriched in clusters 8 and 3 and zones 6 and 7 of the endostyle (Jiang et al., 2023). *Muc5a* and *VWF* transcripts were found in Clusters 3 and 10(Sasaki et al., 2003; Yamagishi et al., 2022). Glutathione peroxidase, a common endostyle and branchial sac enzyme (Kobayashi et al., 1983), was observed in our dataset in Clusters 3 and 6 (Fig. 5A).

**Figure 5.**
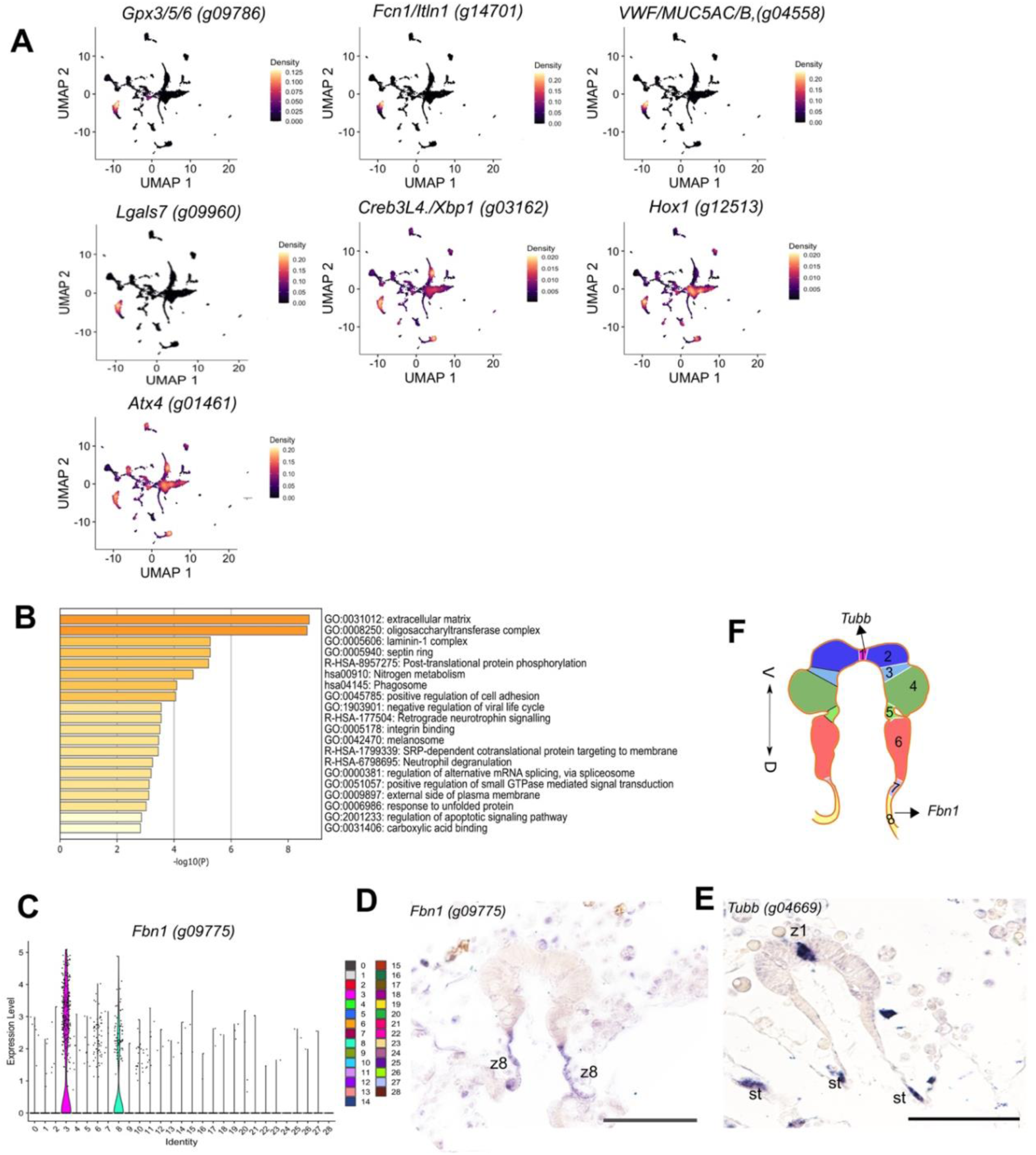
Identification of Endostyle Clusters. A. UMAP feature density plots showing the highest concentration of cells expressing each of these tunicate endostyle genes. B. GO and pathway terms colored by p-value as determined by Metascape. C. UMAP plot for cluster marker 3, *Fbn1*. D. *In situ* expression pattern for Cluster 3 marker *Fbn1*, in zone 8 of the endostyle. E. *Tubb* mRNA was detected in zone 1 cells of the endostyle. F. Schematic showing the different zones of the endostyle, highlighting the spatial expression of *Fbn1* and *Tubb*. Abbreviations: zone 8 (z8), zone 1 (z1), stigmata (st), ventral (v), dorsal (d). Scale bar is 50 μm.

GO and pathway analyses for Cluster 3 associated genes revealed enrichment for processes such as immunity, ECM, carbohydrate binding, and cell adhesion (Fig 5B). Jiang et al. (2023) also found a list of similar GO terms including ribosomal, thyroid hormone, immune function, digestive function (such as mucus), and neurosecretory genes (such as semaphorin 1a) for *Styela clava* (Jiang et al., 2023).

The top marker gene for Cluster 3, g09775, (Fig. 5C) encodes fibrillin (Fbn1), a protein with fibrillin repeats and an EGF domain that shares 44% identity with the Human Fbn1/2/3 proteins. Fbn proteins are found in connective tissues and maintain the elasticity of the tissues. It is critical for microfibril synthesis, as it binds to calcium and regulates TGF-β release, growth, and development (Chaudhry et al., 2007; Handford, 2000; Reinhardt et al., 1996). *Fbn1* expression was observed in zone 8 of the endostyle (Fig. 5D). This region of the endostyle functions with zone 7 in immune activities, with high iodine and peroxidase activities (Jiang et al., 2023; Sasaki et al., 2003) (Alesci et al., 2022). Iodine metabolism has been linked to thyroid gland evolution (Fujita and Nanba, 1971). We also found that the *Tubb* probe (identified as a ciliated cell marker in Fig. 3D) marked a subset of endostyle cells. *In situ* hybridization showed that its mRNA was present in zone 1 cells of the endostyle (Fig. 5E). These cells have long cilia, which together with mucus, aid in trapping food particles from the water current (Holley, 1986). Based on this information, Cluster 3 was assigned as an endostyle cell cluster.

### Identification of Blood cell clusters

The top Cluster 25 gene g07537 is predicted to encode a FAD-dependent oxidoreductase domain-containing protein (Foxred2) (Didion et al., 2002). This transcript was also detected in some of the cluster 12 cells (Fig. 6A). Foxred2 functions to balance redox states and contributes to the generation of reactive oxygen species (ROS), which are associated with endoplasmic reticulum (ER) stress. Strong Foxred2 staining was observed in the thin endothelium lining blood vessels and in a small number of immunocytes with a few large granules (Fig. 6A). Additionally, staining was observed in the developing heart tube cells as they became thinner. In ascidians, the expression of Foxred2-like proteins in the vascular lining and granular immunocytes suggests that it may be crucial for maintaining redox balance and metabolic processes critical for vascular function and immune response. This may include protecting cells from oxidative stress, aiding in antimicrobial defence, detoxifying harmful substances, and regulating inflammation. GO analysis (Fig. S10) revealed associations with heme binding, biosynthesis, oxidoreductase activity, and SLC-mediated transmembrane transport within Cluster 25. SLC-mediated transport is important for thin-monolayer barriers, such as the blood-brain barrier, to mediate the transport of substances across the endothelium (Morris et al., 2017).

**Figure 6.**
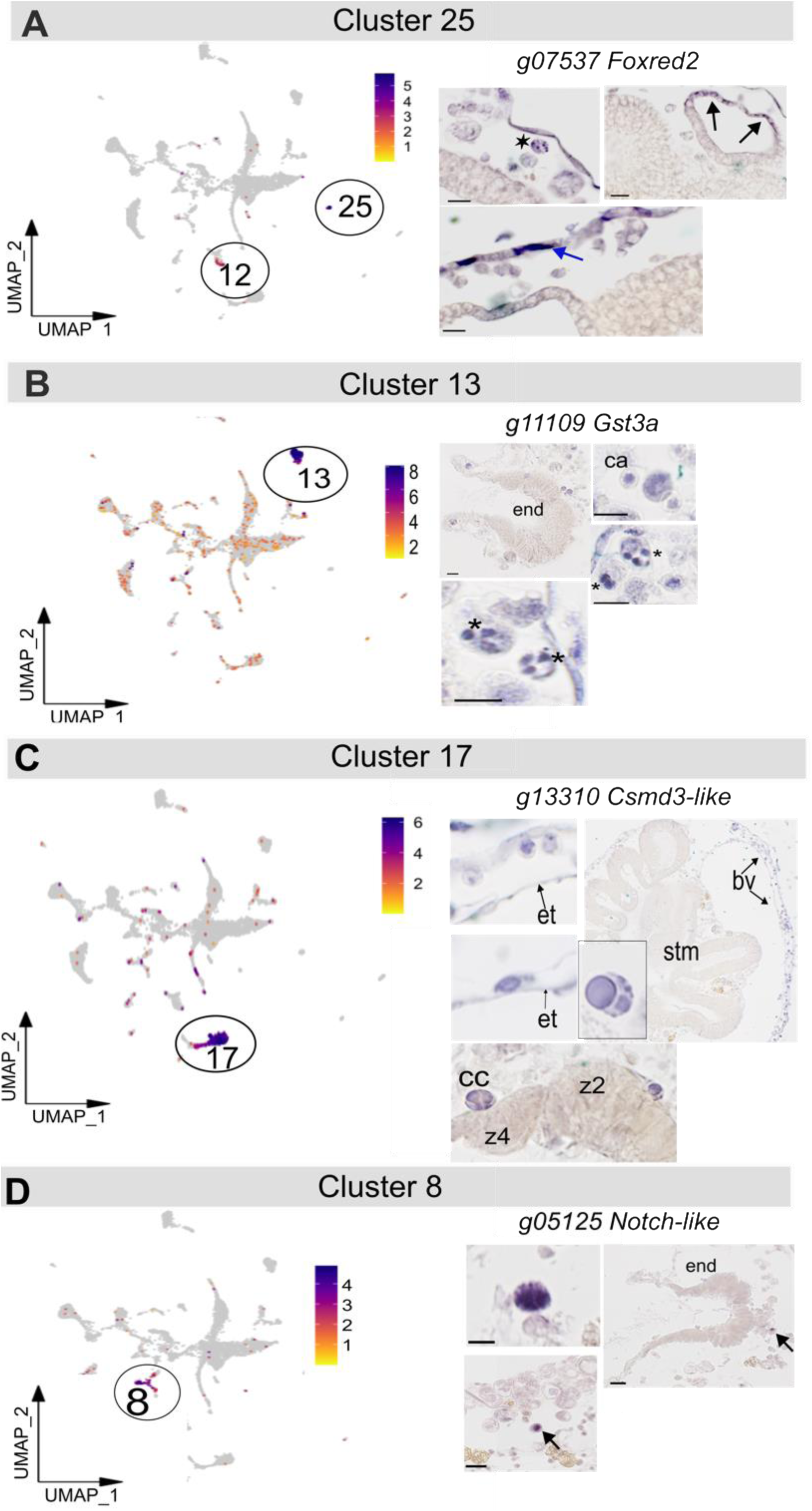
Candidate blood cell clusters. A. UMAP feature plot for Cluster 25 marker, *Foxred2*. Detection of Foxred2 mRNA by in situ hybridisation identified vascular endothelium (blue arrow), cells of the developing heart and some blood cells (asterisk). B. Cluster 13, g11109 *Gst3a*. C. Cluster 17 marker, g13310 (*Csmd3*). *In situ* hybridization with *Csmd3,* with mRNA-positive vesicles indicated (asterisks). D. Cluster 8 marker *g05125* (*Notch-like*). Only a few scattered cells in the vascular stained with the *g05125* probe (arrows). Abbreviations: stomach (stm), vessel endothelium (et), blood vessel (bv), endostyle (end), compartment amoebocyte (ca), compartment cell (cc), zone 2 (z2), zone 4 (z4) The scale bars are 10 μm.

The expression of the Cluster 13 marker, Glutathione S-Transferase Alpha 3 (*Gst3A*) *g11109,* was also detected in some cells and distributed throughout the other clusters (Fig 6B). Staining was observed in cytotoxic cells, including granular amoebocytes (Fig.6B, asterisks), compartment amoebocytes, and morula-like cells (Fig. 6B). Morula cells are variable in size, and the extent of staining may have been influenced by vacuole size. Only a few significantly enriched GO terms were identified because of the low number of marker genes in this cluster. However, it included immune system processes (Fig. S10).

The gene g13310 was identified as a cell marker for Cluster 7 (Fig. 6C). This gene is predicted to encode a protein ortholog of CSMD3 with multiple Sushi and von Willebrand factor type A (CUB) domains, typically found in transmembrane receptors or adhesion proteins. Sushi or complement control domains are involved in the immune system (Ermis Akyuz and Bell, 2022). Excessive activation of the complement pathway is prevented by Sushi domain-containing proteins, which bind to activated C3/4 components to target them for degradation (Ojha et al., 2019). *Csmd3+* cells were detected in the circulation (Fig. 6C), which appeared to be phagocytic cells, including macrophage-like cells and hyaline amoebocytes. Gene ontology analysis revealed an overrepresentation of terms related to the cytoskeleton, apoptosis, and immunity (Fig. S9).

Cluster 8 top marker gene, *g05125* (Fig. 6D), codes for a large protein that shares 30% identity with NOTCH1, NOTCH2, and SNED1 due to multiple Sushi and EGF domains. Several Notch-like genes are present in the *B. diegensis* genome. *In situ* hybridization detected expression in a small number of storage and/or granular cells; larger cells were characterized by multiple small vesicles (Fig. 6D). These cells resemble mast cells and are predicted to release inflammatory factors such as histamine and chemokines (Blanchoud et al., 2017; Cima et al., 2001). Gene Ontology and pathway analysis indicated terms related to peroxidase and leukocyte-mediated immunity (Fig. S11).

### Developmental Trajectory analysis

To identify the cells forming the bud disc of the peribranchial epithelium, we examined cells with thickened epithelium (Berrill, 1947b) and budlet/vesicle precursors. We focused on identifying the cells with the highest expression of transcription factors found in disc cells, *Pitx1, Otx, Nk4*, and *Runx* (Langenbacher et al., 2015; Ricci et al., 2016; Tiozzo et al., 2005). The subset of cells within Cluster 6, likely the peribranchial cluster, showed the highest joint density, indicating that cells co-expressed these four genes (Fig. 7C). Using this as the root, the predicted cell trajectory was plotted using the Monocle3. Pseudotime trajectory paths indicate the progression and pathways of these cells, going from this putative stem cell cluster to more differentiated states later in the pseudotime. This analysis connected all Seurat clusters, with the origin located in Cluster 6 (Fig. 7C). To extend this further, we determined whether several previously studied candidate stem cell markers were present in the scRNA-seq dataset, including *Itga6*, *Notch2*, *Vasa*, and *Piwi* (Kassmer et al., 2020; Kawamura and Sunanaga, 2010; Rinkevich et al., 2010; Rosental et al., 2018). *Vasa, Piwi1*, and *Piwi2* were missing from the dataset. *Notch2* and *Itga6* were detected in the cells scattered across several clusters (Fig. S12). The proportion of dense cells was the same as that of bud disc genes (Fig. S12 and Fig. 7). Additionally, orthologs of the reprogramming factors (Yamanaka factors) (Takahashi and Yamanaka, 2006, 2016), *Oct3/4 (*POU3/4 proteins), *Sox2* (SOXB subgroup of Sox factors), *Klf4*, and *cMyc* were found in the same group of cells within cluster 6 (Fig. S13). This further supports the notion that a subset of Cluster 6 cells likely has stem cell properties and represents the bud disc.

**Figure 7.**
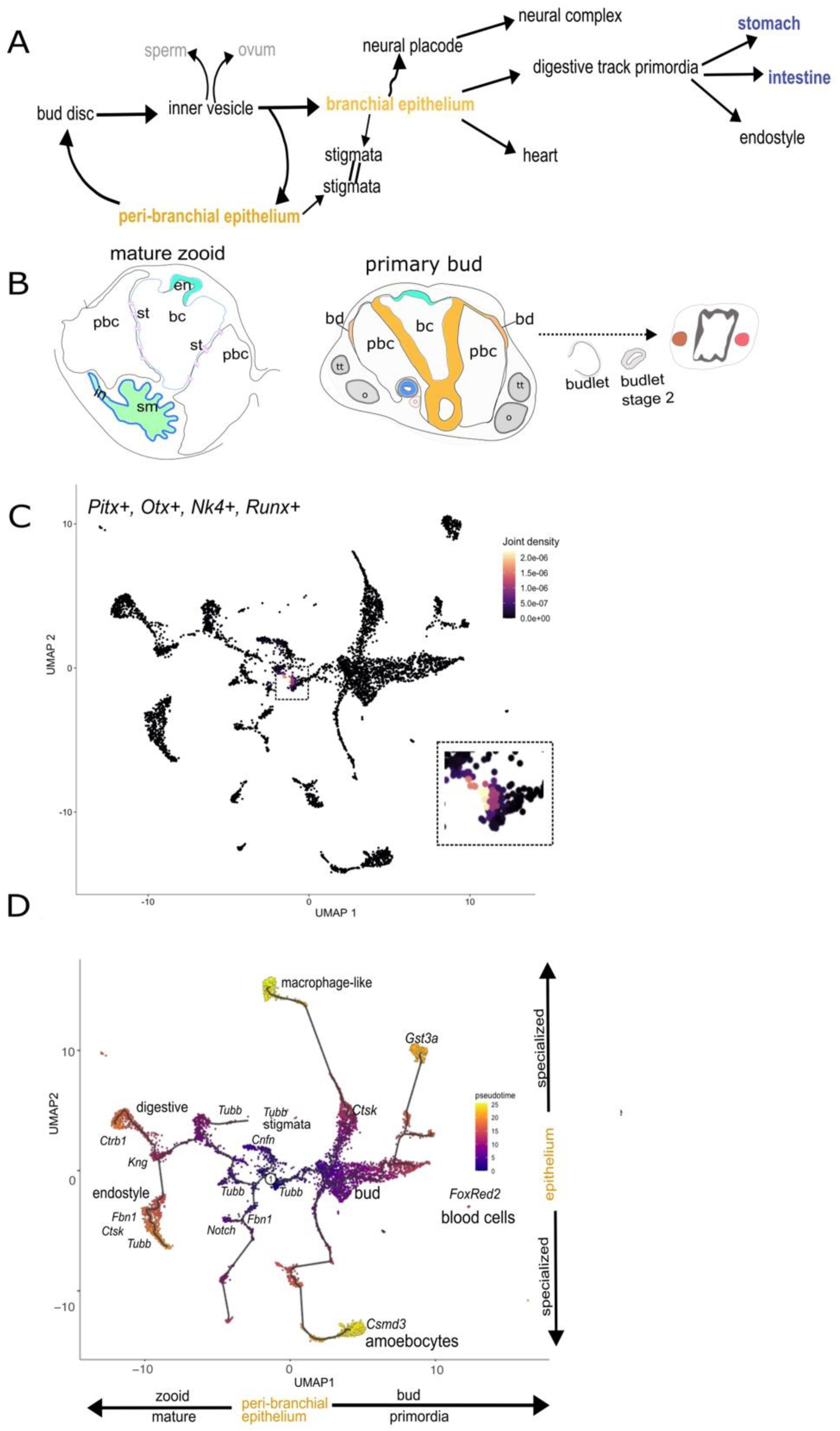
Cell populations and their relationships. A. Summary of the main tissue and organs formed. B. Example schema showing the relationships between tissues. The bud arises from a small group of cells (3-4 cells) within the epithelium of the peribranchial chambers, which appear in the immature zooid before the stigmata perforate. These cells form the bud disc. (Berrill, 1947). The disc separates from the epithelium to form a closed vesicle surrounded by the outer epidermis obtained from the parent zooid. Germ cells segregate early from the inner vesicle and form oocytes and sperm. On the dorsal side of the vesicle, the invagination forms a neural placode. The heart and digestive tracks form on the posterior side, and primordia form first as outpockets of the branchial epithelium. C. UMAP density plot showing co-expression of *Pitx, Otx, Nk4*, and *Runx* in cells located in Cluster 6. Cells with higher joint densities were more orange in color. D. Monocle 3 trajectory analysis. The root is the progenitor (within cluster 6, likely the peribranchial epithelium). Abbreviations: endostyle (en), stigmata (st), peribranchial chamber (pbc), branchial chamber (bc), intestine (in), stomach (sm), testes (tt), ovary (o), bud disc (bd).

The trajectory of the colony dataset was analyzed to understand the relationship between cell clusters and differentiation time using Monocle3 (Fig. 7D). The trajectory of cell development was charted on UMAP, with the candidate progenitor bud disc cells of the peribranchial cluster (Cluster 6) serving as the root (Fig. 7C). Pseudotime trajectories reveal that the clusters, predicted to be bud, peribranchial and brachial epithelial cells, emerge early. In contrast, clusters associated with specialized tissues, such as the stomach epithelium and neuronal clusters, were later in the pseudotime (Fig. 7D). Epithelial tissues within the zooid appeared early in the trajectory, preceding the development of stomach and endostyle tissues (left branch). The third pathway leads to immune cell formation. Overall, the development of zooid-related tissues later in the trajectory was consistent with blastogenesis (Fig. 7A and B)

To explore the relationships between clusters, we generated PAGA plots to investigate the underlying structure and infer the potential differentiation pathways (Fig. 8A). This suggests that multiple common progenitor populations may give rise to differentiated cell types. The relationships between different clusters of cells during colony asexual reproduction were visualized, with each node representing a distinct cluster identity. Lines connecting the nodes illustrate lineage relationships between clusters. By examining central nodes with numerous connections, such as nodes 20, 21, 0, and 6, we can identify potential progenitor states that can differentiate into several other states. These clusters had numerous outgoing edges, indicating potential branching points in the differentiation trajectories (Fig. 8A). Finally, the cell clusters at the ends of the branches represent the terminally differentiated states, and this largely matches the later cell Clusters identified in pseudotime analyses (Fig. 8C and 7D).

**Figure 8.**
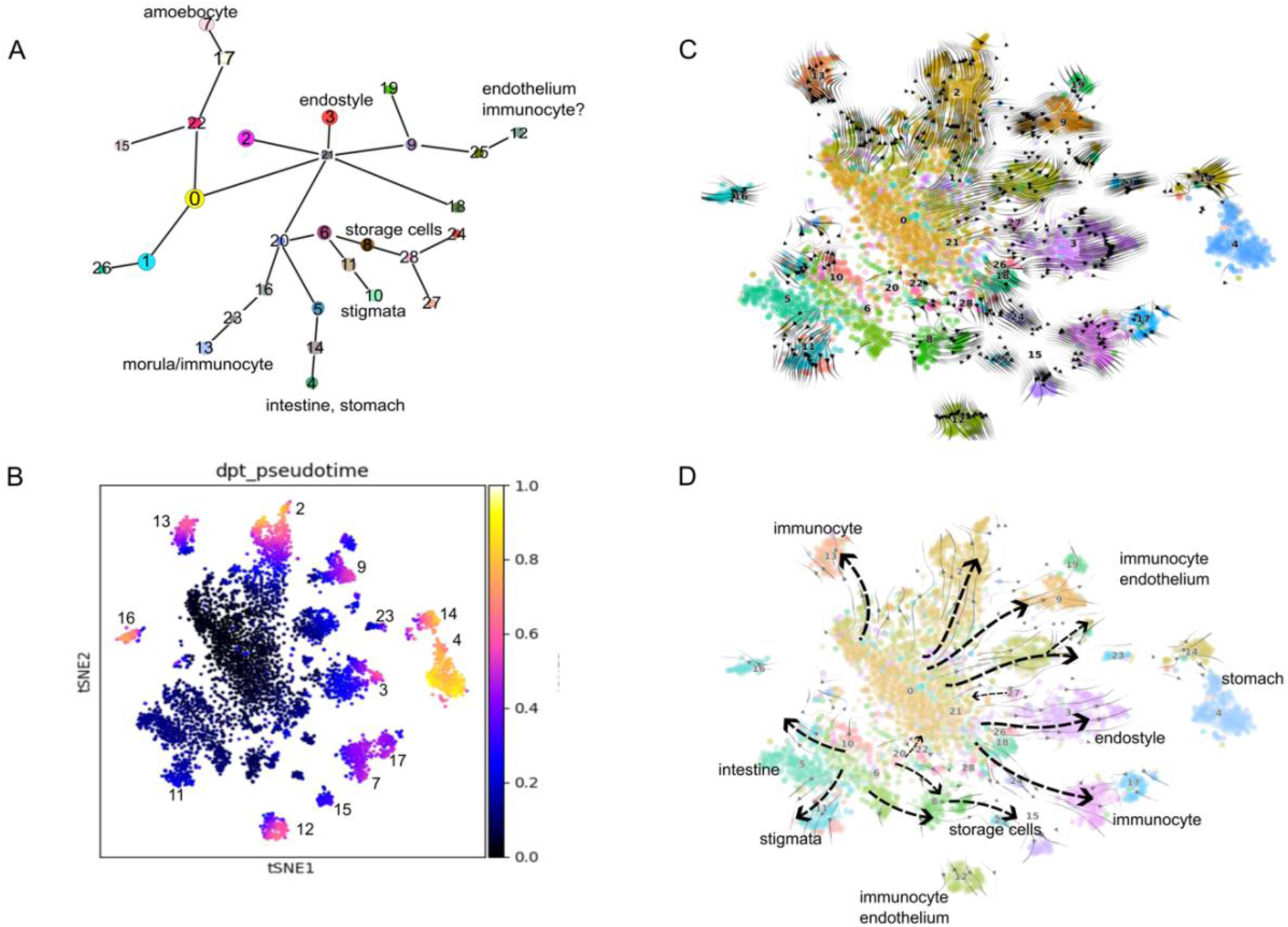
Identification of potential differentiation pathways using CellRank. A. The PAGA plot displays each cell cluster as a node, with branching lines indicating potential differentiation pathways among the clusters. B. A tSNE projection where cells are colored by pseudotime, showing the progression of cell states over time. Darker colors represent earlier pseudotime states, whereas lighter colors represent later pseudotime states. The Cluster IDs are given for the later states. A UMAP projeciont is shown in Fig. S15. C and D. Velocity plots using the tSNE projection, where arrows represent the direction of RNA velocity, indicating the predicted future states of the cells based on their transcriptional activity and pseudotime. The cells are colored using the Seurat Cluster ID. D. An annotated version of plot C shows detailed identification of specific cell types and their predicted differentiation pathways.

The velocity (scvelo) plot utilizes the spliced-to-unspliced mRNA ratio to indicate the direction of the cell transition. The length of the arrows in the velocity plot represents the speed of the transitions, with longer arrows indicating faster transitions and shorter arrows indicating slower or more stable transitions. This provides dynamic information about cellular processes. Additionally, the arrows converged at specific points, offering further insights into the directionality and progression of cell states. Clusters with numerous outgoing arrows are highlighted, emphasizing their roles in cellular differentiation (Fig. 8C). This plot was consistent with the PAGA plot, particularly with multiple transition events from clusters 0, 6, 21, and 20 (Fig. 8A), suggesting that cells within these clusters serve as central nodes with multiple connections (Fig. 8D).

Finally, CellRank was used to examine the cell fate dynamics (Fig. 9). It has been designed to deal with complex data, such as data produced from developing and regeneration tissues, with many initial, intermediate, and terminal cell states (Lange et al., 2022). Figures 9C and S14 display a bar graph of the aggregate fate probabilities of cells from the Seurat cluster. These cells will progress to a terminal state, and the clusters with multiple bars of varying height have different probabilities of transitioning to different end states, such as Clusters 0, 5, 6, 8, 20, 18, 21, 22, 26, and 28. In contrast, clusters with cells that transition to one fate predominantly have a single high bar. For example, all cells in Cluster 11 transitioned to the ciliated stigmata cell type, according to GO and *in situ* expression (Fig. 3C).

**Fig. 9.**
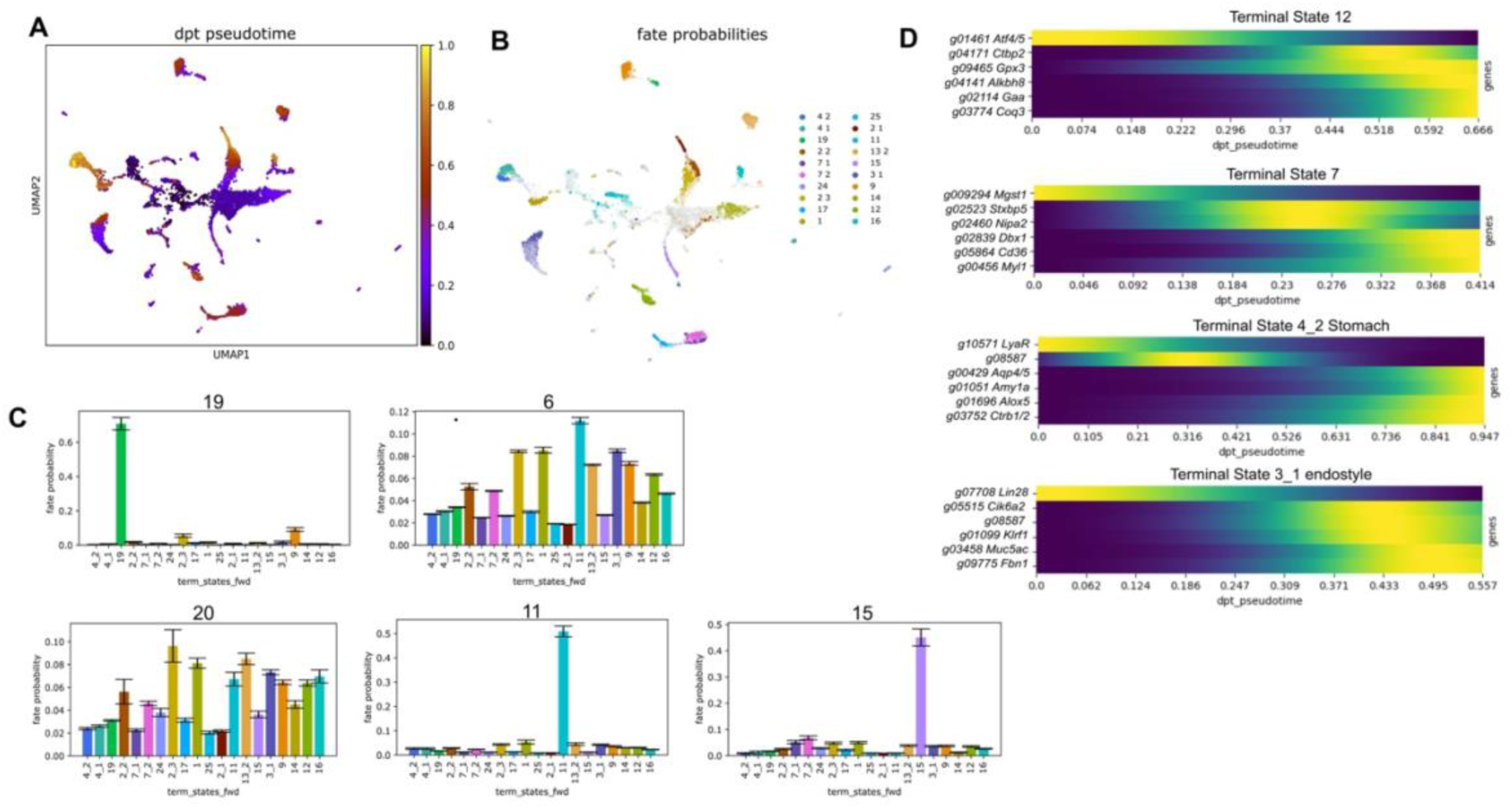
Cell fate dynamics. A. UMAP plot showing diffusion pseudotime (dpt_pseudotime). The color scale represents pseudotime, cells colored yellow appear later in the predicted cell trajectory pathway. B. Fate probability UMAP, shown as terminal clusters. C. Aggregate fate probabilities fate bar graphs. These display the probabilities of different clusters (22, 11, 15, 19, and 6) progressing towards the terminal states, as predicted by CellRank. D. A selection of driver genes is displayed alongside a gene whose expression is highest at the earliest pseudotime on the lineage towards the terminal state fate (File S4).

We used CellRank to identify potential driver genes strongly associated with the terminal state that may serve as key regulators or markers of cell fate (Fig. 9E and File 4). These genes exhibited high expression levels at the earliest pseudotime but were low in the final terminal state. Such genes may play a role in initiating or driving the early stages of differentiation and then decreasing their expression as cells reach their final differentiated state. One example of a potential driver gene is the *Lin2*8 ortholog, a highly conserved RNA-binding protein that regulates stem cell differentiation and proliferation in organisms ranging from nematodes to mammals (Wu et al., 2022). This gene is expressed in undifferentiated pluripotent cells and is downregulated as cells progress toward a specialized cell fate or undergo reprogramming (Pieknell et al., 2022). Another potential driver gene was *the Atf4/*5 ortholog. In vertebrates, this transcription factor is associated with stress-induced responses and the differentiation of stem cells into secretory cells. It plays a role in epithelial differentiation and stresses immune response (Barrera-Lopez et al. 2024). *B. diegensis g11626* encodes a DExD/H-box polypeptide, 39 B (DDx39b)-like RNA helicase. DDx39b regulates mRNA splicing events, including the processing of transcripts of genes required for immune and myocyte cell fate (Hirano et al., 2023; Zhang et al., 2018). Another potential driver gene is high mobility group box 1 (*Hmgb1*), a chromatin-associated protein secreted by immune cells and involved in differentiating mesenchymal stem cells into vascular cells (Meng et al., 2018).

Together, these analyses provided strong supporting evidence for the inferred differentiation pathways. They identified key transitional cell clusters and several candidate progenitor cell populations.

## Discussion

Using sc-RNA-seq, we gained insights into the cellular content of a *B. diegensis* colony. This technology identifies cluster markers by deconstructing the cellular heterogeneity of the filter-feeding colony tissues. The colonial tissue was resolved into 29 clusters, including zooid and vascular tissues. As a more ancestral chordate, *B. diegensis* occupies an essential phylogenetic space with vertebrates, highlighting its significance in evolutionary and developmental research. This sc-RNAseq resource paves the way for examining the cellular composition of the colonial chordate, *B. diegensis*.

Based on the marker expression profiles that were highly and distinctively expressed in each cluster, the colonial cellular profiles showed conserved gene regulatory programs and functions. In contrast, clusters not enriched for a particular group of markers or functional annotations may indicate an intermediate cell state. Although we found some blood cell markers, some predicted hemocyte clusters showed intermediate and non-specific gene expression patterns. These functions were not specifically enriched for the blood type because of functional annotations based on the closest vertebrate relatives. Similarly, some blood populations of *B. schlosseri* did not exhibit specific functions and contained unannotated genes with invertebrate homologues (Rosental et al., 2018).

Trajectory analysis of cell fate during blastogenesis suggested that multiple initial and intermediate states are present in the colony at any given time. In ascidians, bud development is initiated by the emergence of bud primordium through the thickening of the peribranchial epithelium via dedifferentiation or transdifferentiation (Manni et al., 2007). The peribranchial epithelium corresponded to Cluster 6 in our study. Our gene expression analysis in Cluster 6 supports the hypothesis that the peribranchial epithelium is a critical source of progenitor cells for bud development. The endostyle has been proposed as a stem cell niche in zooids, suggesting a more transitional stem cell source for the colony (Voskoboynik et al., 2008a). Specialized progenitors associated with the vascular lining and epithelial tissues have also been identified. This demonstrates the ability of epithelial tissues to function as a source of progenitors, preserving progenitor-like characteristics. This is significant because it indicates that *Botrylloides* stem cells may not be fixed in one location, but can transition between different niches within the organism. Additionally, the presence of transitional stem cell niches, such as endostyle, highlights the dynamic nature of stem cell populations in these organisms.

Adult stem cells (ASCs) are involved in various epithelial tissues in invertebrates. For example, in sponges, epithelial cells function similarly to stem cells and play a role in tissue repair (Rinkevich et al., 2022). This observation suggests a conserved mechanism across different invertebrate species where epithelial cells have multipotent capabilities essential for regeneration and repair. This comparison underscores the importance of epithelial and vascular progenitors in colonial ascidians and other invertebrates, indicating a potentially conserved evolutionary strategy for tissue regeneration and maintenance. We hypothesized that epithelial-and vascular-associated progenitors play key roles in the development and maintenance of colonial ascidians.

Assigning cellular identities in scRNA-seq is inherently challenging because of the complexity and variability of the gene expression profiles. The study was based on a limited set of approximately 6,000 cells, which may not capture the full diversity of cell types in the organism. Low-abundance transcripts may be underrepresented, leading to a potential bias in gene expression data and overlooking genes expressed at low levels. Obtaining cells from the tunic, which is rich in cellulose, presents a significant challenge, leading to the absence of certain tunic cell types. The timing of colony collection might have missed some stages of the sexual cycle, potentially omitting important cell types or developmental stages. For example, the stages of oogenesis will be missed as *B. diegensis* only sexually reproduces briefly at the end of the summer in New Zealand.

*In situ* hybridization, which is useful for the spatial localization of gene expression, is difficult to quantify accurately, limiting its utility in confirming single-cell RNA sequencing (scRNA-seq) findings. No one cluster was identified as marked by gene expression commonly attributed to stem-like cells in colonial ascidians (Ballarin and Rosner, 2022). Specifically, previously identified germline markers for colonial and solitary tunicates, such as *Piwi* or *Vasa* (Kawamura et al., 2011; Rinkevich et al., 2010), were missing from the dataset. This absence could be due to technical dropout, a common issue in scRNA-seq where lowly expressed genes might not be detected in every cell, leading to their apparent absence in the data (Kharchenko et al., 2014). The genes in *B. diegensis* were named and assigned functions for gene ontology analysis based on their closest vertebrate orthologs, assuming similar functions in ascidians. However, many genes lack orthologs, making functional prediction more challenging.

## SUMMARY

In this study, we performed single-cell RNA sequencing (scRNA-seq) on an entire *B. diegensis* colony, including zooids, buds, and the vascular tunic, to identify cell and tissue markers and resolve cellular heterogeneity. The analysis identified 29 major cell clusters within the colony, and *in situ* hybridization was used to examine the spatial expression of the cluster marker genes. Various tissue types have been identified at the molecular level, including blood cells and zooid tissues such as the branchial epithelium, stomach, and endostyle. Distinct cluster markers were found in specific regions of the stomach epithelium, highlighting their specialization. Trajectory estimations were aligned with predictions based on the early appearance of progenitor clusters, whereas zooid-related tissues appeared later in the developmental path. This study provides a valuable resource for understanding the development, function, and regeneration of *B. diegensis*, and demonstrates the power of scRNA-seq in defining cell types and tissues in complex colonial organisms.

## Methods

### Animal husbandry

Colonies of *B. diegensis* were sampled from Otago Harbor in New Zealand (45°52’18.1″’’S, 170°31’37.6’’″ E). The animals were attached to 5x7 cm glass slides. The tanks were aerated, and the colonies were fed a shellfish diet (a blend of marine microalgae). The colonies were fed every two days, and the seawater was replaced. Colonies were confirmed to be *B. diegensis* by COI barcoding (Temiz et al., 2023).

### Single-cell preparation & FACS

Single cells of *B. diegensis* were prepared using the ACME dissociation method described by García-Castro et al. (2021). Fresh ACME solution was prepared for fixation and separation of tissue cells. It contained DNase/RNase-free distilled water, methanol, glacial acetic acid, and glycerol at a ratio of 13:3:2:2. The mature colony was in stage A of blastogenesis with active filtering, newly emerged primary buds, and no signs of secondary buds. The colony was placed in a Petri dish on a glass slide, where it was attached using a microtome razor blade with minimal disturbance. The animal was washed with 1 ml of 7.5% N-acetyl l-cysteine in 1XPBS. This solution removes excess seawater and protects RNA. One mL ACME was added, and the colony was minced well using a single-edged razor blade.

The suspension and all the larger tissue pieces were placed in a 1.5 mL tube. The tube was placed in a rotator to apply seesaw motion at ∼30-40 rpm for 1 h at room temperature. The sample was pipetted twice and strained with a 40 µm cell strainer into an ice-covered 50 mL Falcon tube. The cells were cooled to prevent RNA degradation. The suspension was centrifuged at 1000 *g* for 5 min at 4°C. The supernatant was removed, and the pellet was apparent. One mL of 1% BSA-1XPBS with RNase inhibitor was added to the pellet, and the tube was flicked to mix. As the second washing step to remove all ACME, 1 mL of 1% BSA-1XPBS with RNase inhibitor was added to the tube, and the tubes were flicked. A 70 µm flowmi cell strainer was used to strain the cells to decrease the number of aggregates. The sample was centrifuged at 1500 *× g* for 5 min at 4°C, and the cell pellet was visible.

The cells were checked on a hemocytometer by staining with trypan blue (1:1). The approximate number of cells was estimated, and their integrity was investigated. ACME fixed cells were stained with the DNA dye, DRAQ5^TM^, to sort intact single cells from cellular debris and aggregates (eBioscience 0.66 μL/mL of 5 mM stock). After staining, cells were incubated in the dark and on ice for an hour. The stained cells were sorted using a BD FACSAria Fusion flow cytometer (BD Biosciences) with a red laser (640 nm). In total, 50,000 cells were sorted in collection buffer containing 1X PBS-1% BSA-RNAse inhibitor (40 U/mL) (Fig. S1). The cells were visualized under the far-red channel of a Nikon Ti2 Inverted fluorescence microscope (Fig. S1). The cells were preserved at -20°C by adding DMSO (10% final concentration).

### sc-RNA-seq via 10X Genomics & sequencing

Frozen cells were thawed on ice and centrifuged at 1500 × g and 4°C. The supernatant was used to wash away the DMSO, followed by adding 1XPBS-1%BSA-RNase inhibitor (40 U/mL). Sorted cells were counted on a hemocytometer using trypan blue in a 1:1 ratio and diluted to yield approximately 6000 cells for further processing. The cells underwent centrifugation at 1500 × g at 4°C, then 10X Master Mix was added to the single cells. Subsequently, the cells were loaded onto a 10x Genomics Chromium chip. Gel bead emulsion (GEM) generation, barcoding, reverse transcription, cDNA amplification, and library preparation were carried out according to the 10X Genomics protocol for 3’ Gene Expression (v3) user guide (10X Genomics CG000183 Rev C). The target cell recovery rate was set to 3000. After the GEM generation, a post-GEM-RT clean-up procedure was performed. cDNA amplification was performed for 16 cycles. The resulting cDNA was analyzed using a Qubit fluorometer and agarose gel electrophoresis. Samples were indexed for library preparation and library concentrations were quantified using KAPA PCR. The combined cDNA libraries were sequenced with a length of 150 bp on an Illumina NovaSeq 6000 sequencing platform at the Australian Genome Research Facility Ltd.

### Read alignment & cluster analysis

Quality checks of the library were performed using FastQC (Andrews, 2010) (Supplementary Fig. S22). Raw sequencing reads were mapped to the reference *B. diegensis* genome using STARSolo (Kaminow et al., 2021). The genome file of *B. diegensis* (formerly *B. leachii*) was downloaded from the Aniseed Database (http://www.aniseed.cnrs.fr) (Blanchoud et al., 2018). Gene models were identified using the StringTie software (Pertea et al., 2016). First, genome indices were created using STAR 2.7.9 (--genome SAindexNbases 12)(Dobin et al., 2013). The indices were then used to map the raw sequences to the *B. diegensis* genome using STAR 2.7.9, which masks the polyA tail during alignment; therefore, no prior trimming was performed. These run options were selected for the default barcode lengths using a droplet-type algorithm (soloUMIlen 12, soloType Droplet). No barcode read length defined (-- soloBarcodeReadLength 0). The empty droplets were then filtered (soloCellFilter EmptyDrops_CR). Finally, sequences with barcodes present within the barcode whitelist were selected while mapping (soloCBwhitelist). The mapping statistics are listed in Table S1.

After mapping, clustering was performed using Seurat 4.0.1(Stuart et al., 2019) and R 4.1.3. Before clustering, filtering was performed by selecting transcriptomes with 200–2000 genes expressed in at least three cells. The counts were normalized to the total counts using log normalization, and the scale factor was set to 10000. Variability was identified within the 2000 genes using the *FindVariableFeatures* function with vst as the selection method. The linear dimensional reduction method was applied to the single-cell transcriptome using principal component analysis. The first 50 PCs for the mature colony were selected based on elbow and jackstraw analyses. Clustering was executed with a 0.8 resolution determined using the clustree R package (Zappia and Oshlack, 2018). The data were plotted using the nonlinear dimensional reduction method and presented using the Uniform Manifold Approximation and Projection (UMAP).

Differential expression analysis was performed using Seurat package to identify the top genes that were potential markers for each cluster. Top genes were considered significant if Padj < 0.05 and had a log_2_FC (fold change) value of > 0.5 (Supplementary File 1). Single-cell transcriptomes were saved in the Seurat object, which can be used to check for feature expression. Finally, gene annotation was performed using Metascape (Zhou et al., 2019). The pathway and GO term enrichment were calculated by comparing them to the background genes (all the expressed genes in the dataset with an orthologue, 6039 genes).

Pseudo-time estimations of single cells were calculated using Monocle 3 (Trapnell et al., 2014), Scanpy 1.10.1 (Wolf et al., 2018), scVelo 0.3.2 (Bergen et al., 2020) and CellRank v. 2.0 (Lange et al., 2022). Code availability: https://github.com/MJWilsonOtago/scRNAseqBotrylloides/.

### Probe synthesis*, in situ* hybridization & image acquisition

Probe preparation was initiated by cloning the target genes. Primers were designed to amplify the genes of interest (Table S2). The amplified fragments were inserted into pCRII-TOPO (Life Technologies). Plasmids were sequenced to verify the correct fragment. Digoxygenin (DIG)-labelled sense and antisense probes were synthesized using a 10× DIG RNA labelling mix and SP6/T7 RNA polymerases (Sigma-Aldrich) via *in vitro* transcription. In situ hybridisation was performed as described previously (Zondag et al., 2019). Images were acquired using a Nikon TiE with 60x magnification.

## Supporting information

Supplementary Tabes

## Data availability

The authors declare that all data supporting this study’s findings are available in the article and its supplementary information files or through https://megan-wilson-otago.shinyapps.io/Botryllodies_scRNAseq/.

## Acknowledgements

The authors would like to thank the Gemmell Lab of the University of Otago, particularly Joanne Gillum, for their contribution to the single-cell optimization. We also thank Bridget Fellows, Devon Gamble, Justine Gapuz, and Ed Moody for their comments on the final draft. This study was supported by funding from the Department of Anatomy and a University of Otago Research Grant. B.T. was supported by an Anatomy Department Ph.D. scholarship from the University of Otago.

## Contributions

B.T.: Data acquisition and interpretation, and manuscript writing. M.M.: funding acquisition, data acquisition, interpretation, and manuscript writing. M.J.W.: Study design, data acquisition and interpretation, funding acquisition, and manuscript writing.

