## Supplementary Tabes for "Single-cell transcriptomic profiling of the whole colony of *Botrylloides diegensis*: Insights into tissue specialization and blastogenesis"

### Supplementary Tables

**Table S1.** STARSolo mapping summary stats for the whole colony.

| **Mapping Summary** | **adult** |
|---|---|
| Number of Reads | 57912212 |
| Reads With Valid Barcodes | 0.966289 |
| Sequencing Saturation | 0.703259 |
| Q30 Bases in CB+UMI | 0.951488 |
| Q30 Bases in RNA read | 0.949242 |
| Reads Mapped to Genome: Unique+Multiple | 0.824445 |
| Reads Mapped to Genome: Unique | 0.766516 |
| Reads Mapped to Gene: Unique+Multiple Gene | 0.75533 |
| Reads Mapped to Gene: Unique Gene | 0.733161 |
| Estimated Number of Cells | 6353 |
| Unique Reads in Cells Mapped to Gene | 34599629 |
| Fraction of Unique Reads in Cells | 0.814896 |
| Mean Reads per Cell | 5446 |
| Median Reads per Cell | 4058 |
| UMIs in Cells | 10081026 |
| Mean UMI per Cell | 1586 |
| Median UMI per Cell | 1202 |
| Mean Gene per Cell | 481 |
| Median Gene per Cell | 417 |
| Total Gene Detected | 13621 |

**Table S2. Number of DEGs for each single-cell cluster.**

The total number of genes identified is ranked from largest to smallest. The numbers of unmatched genes (lacking vertebrate orthologous) are provided for each

| Cluster | Total genes | Unmatched genes |
|---|---|---|
| Cluster 27 | 479 | 15 |
| Cluster 14 | 465 | 19 |
| Cluster 4 | 405 | 15 |
| Cluster 24 | 393 | 9 |
| Cluster 28 | 381 | 12 |
| Cluster 5 | 378 | 9 |
| Cluster 2 | 374 | 15 |
| Cluster 23 | 370 | 11 |
| Cluster 19 | 364 | 11 |
| Cluster 0 | 350 | 9 |
| Cluster 11 | 338 | 9 |
| Cluster 17 | 337 | 12 |
| Cluster 25 | 317 | 14 |
| Cluster 26 | 284 | 8 |
| Cluster 22 | 283 | 10 |
| Cluster 13 | 281 | 10 |
| Cluster 9 | 277 | 12 |
| Cluster 16 | 267 | 9 |
| Cluster 20 | 225 | 9 |
| Cluster 12 | 215 | 8 |
| Cluster 10 | 214 | 7 |
| Cluster 18 | 210 | 9 |
| Cluster 6 | 206 | 6 |
| Cluster 7 | 198 | 7 |
| Cluster 3 | 187 | 9 |
| Cluster 15 | 182 | 8 |
| Cluster 21 | 166 | 8 |
| Cluster 1 | 163 | 3 |
| Cluster 8 | 147 | 4 |

**Table S3. Number of differentially expressed genes (DEGs) by cluster.**

| Cluster ID | DEGs (> 2-fold enrichment; Padj < 0.05) |
|---|---|
| 14 | 165 |
| 27 | 148 |
| 19 | 112 |
| 24 | 109 |
| 2 | 108 |
| 11 | 88 |
| 4 | 80 |
| 25 | 67 |
| 17 | 58 |
| 10 | 46 |
| 3 | 43 |
| 12 | 43 |
| 4 | 40 |
| 5 | 34 |
| 26 | 30 |
| 28 | 28 |
| 7 | 24 |
| 18 | 23 |
| 15 | 20 |
| 16 | 20 |
| 8 | 20 |
| 13 | 15 |
| 0 | 14 |
| 23 | 13 |
| 6 | 12 |
| 21 | 12 |
| 22 | 11 |
| 1 | 9 |

### Table S4. Genes from the literature and this study that were used support cluster identity.

| Gene ID | Orthologue(s) (Human) | Molecular function | Reference |
|---|---|---|---|
| **Endostyle** | | | |
| g03458 | MUC5AC; MUC5B; VWF | | (Yamagishi et al., 2022) |
| g03162 | CREB3L4; XBP1 | | (Wu et al., 2022) |
| g14701 | FCN1; ITLN1/2 | | (Jiang et al., 2023) |
| g09960 | LGALS7/9 | | (Parrinello et al., 2015) |
| g07754 | FCN2; FIBCD1; MFAP4 | Calcium, carbohydrate binding. Molecular transducer, Lectin signaling | (Jiang et al., 2023) |
| g00874 | LGALS9; LGALS9B/C | Carbohydrate binding, Lectin | (Parrinello et al., 2015) |
| g09483 | PTPRQ | | (Jiang et al., 2023) |
| g09786 | GPX3/5/6 | | (Kobayashi et al., 1983) |
| g01461 | ATF4 | | |
| g09775 | FBN1/2/3 | | This study |
| **Early peribranchial bud** | | | |
| g12344 | RUNX1/2/3 | Transcription factor | (Langenbacher et al., 2015) |
| g08980 | PITX1/2/3 | Transcription factor | (Tiozzo et al., 2005) |
| g09275 | CRX; OTX1/2 | Transcription factor | (Ricci et al., 2016) |
| g13325 | NKX2-3; NKX2-5; NKX2-6 | Transcription factor | (Yamagishi et al., 2022) |
| **Yamanaka factors** | | | |
| g15467 | cMyc | Transcription factor | (Takahashi and Yamanaka, 2016) |
| g11013 | Soxb | Transcription factor | |
| g06244 | Pou3 | Transcription factor | |
| g01331 | KLF5 | Transcription factor | |

#### Table S5. Oligonucleotides were designed to generate *in situ* probes.

| Gene ID | Gene | Primer - Forward | Primer - Reverse | Amplicon Length (bp) |
|---|---|---|---|---|
| g05125 | *Notch1* | GGCTACCAAGGAGATGGATTTAG | TCAGTAAGAGGCGGACATTTG | 683 |
| g13310 | *Csmd3* | GTACACGATCTGCGAAGGAATA | CAACCCGTCAGTCAGGATAAA | 721 |
| g03753 | *Ctrb1* | GGTGTCACTTCGTAGACCTATTG | GCATACACTGCTGGTTGAGTA | 600 |
| g08491 | *Kng1* | AGGCTACTATTTGTGCCTACTTG | AATGGTCTTCCCTTTCGTCTTC | 610 |
| g04846 | *Cnfn* | ACATGGGAGAATGGAATACTGG | ACTAGCGAATTGGTATTCTGTGA | 656 |
| g07537 | *Foxred2* | CAGCGAGGCTTCGGATATTTA | TAACGGGTCGGTAACAAACTAC | 790 |
| g04669 | *Tubb* | GCTGAGGAGGCATAGAGAATAG | GCTGAGACGAGATGGTTAAGA | 866 |
| g04141 | *AlkBh8* | ATCACACGAACGGTGGTAAA | GGGAAGAATGGGAGGGAATATC | 790 |
| g02151 | *Col24a1* | CGCAAGGTAACGAAGGATCA | AATAACCTGGTGGACCCATTAC | 822 |
| g09775 | *Fbn1* | TCGCCTGGAAGTTACGAATG | GTTCCATCCTTGCCCGATAA | 678 |
