## Supplementary Figures for "Single-cell transcriptomic profiling of the whole colony of *Botrylloides diegensis*: Insights into tissue specialization and blastogenesis"

**Supplementary Files**
File 1: Top DEG for each Seurat Cluster, includes genome ID, stringtie_ID and closest vertebrate orthologue.
File 2:
File 3: Results from the GO and pathway analysis with Metascape.
File 4: CellRank driver gene analysis for terminal clusters.

### Supplementary Figures

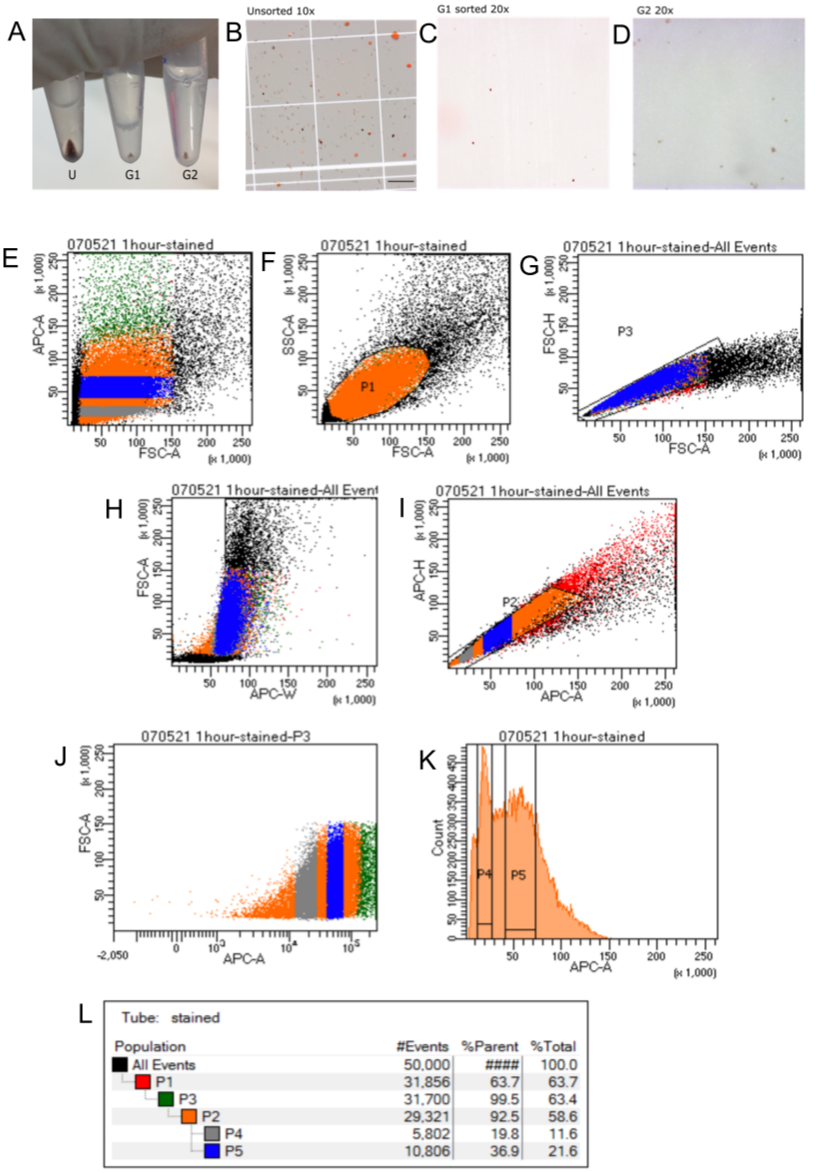


**Figure S1. ACME dissociation and single-cell sorting from mature *B. diegensis* colonies using FACS. A.** The ACME cells were fixed and dissociated. A pellet was visible for the unsorted (U), and sorted G1 and G2 cells. **B.** Single-cells are apparent and intact after ACME fixation observed under fluorescent and bright field microscopy. **C.** G1 cells are present and fluorescent after sorting. **D.** G2 cells were sorted and examined under a fluorescence and bright field microscope. **E.** All events are shown after gating 50,000 single cells stained with DRAQ5. **F.** First gate selection was performed for P1 (population 1) cells based on their ideal area (FSC-A) vs. height (APC-A) signal, because higher and lower ratios are indicative of aggregates and debris, respectively. **G.** Selection of singlets with better height and size ratios within P1. **H.** Width (APC-W) vs. area of the selected P1 and P3 singlets are plotted. **I.** P2 cells were determined based on the most optimal area vs. height ratios. **J.** The area of cells correlated with their fluorescent signals. **K.** Selection of G1 (P4) and G2 (P5) cells based on their area from the P2 cells. **L.** Singlet counts of five populations and percentages of those selected from parent cell populations.

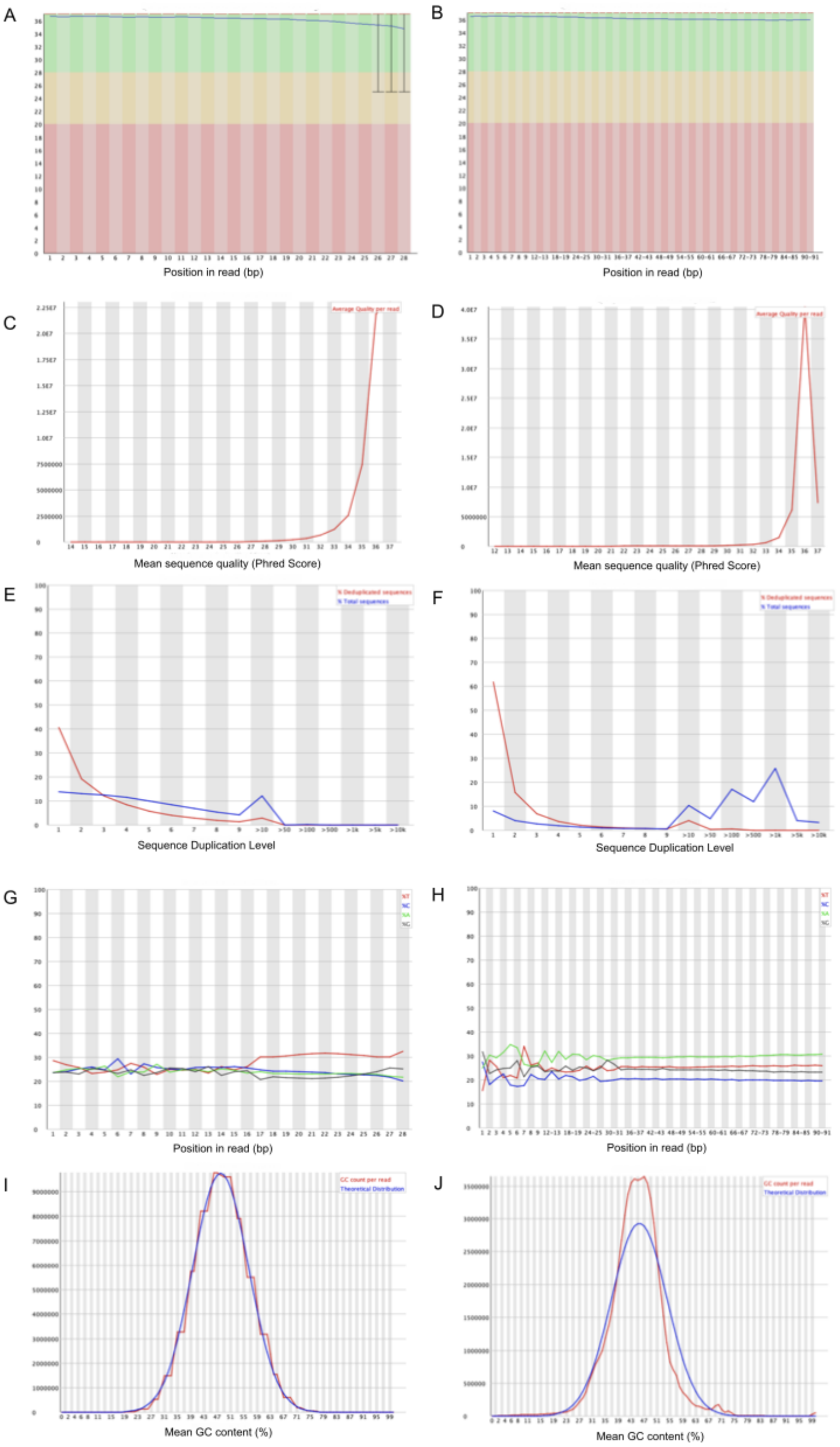


**Figure S2. FastQC quality checks were performed on the library (whole colony).** Quality statistics of paired-end reads are shown for forward reads in A-C-E-G-I and reverse reads in B-D-F-H-J. **A.** Per base quality results showed forward reads with an average size of 28 bp. **B.** Reverse reads per base sequence quality were good, with an average length of 91 bp. **C.** Per-sequence quality scores are shown for forward reads. **D.** Per sequence quality scores are shown for reverse reads. **E.** Duplication of forward sequences was higher than estimated. **F.** Sequence duplication levels for reverse reads are greater than normal. **G.** Per base sequence content for the forward reads is shown. **H.** The sequence content of the reverse reads was plotted. **I.** Normal distribution of GC content for forward reads. **J.** Normal distribution of GC content for reverse reads.

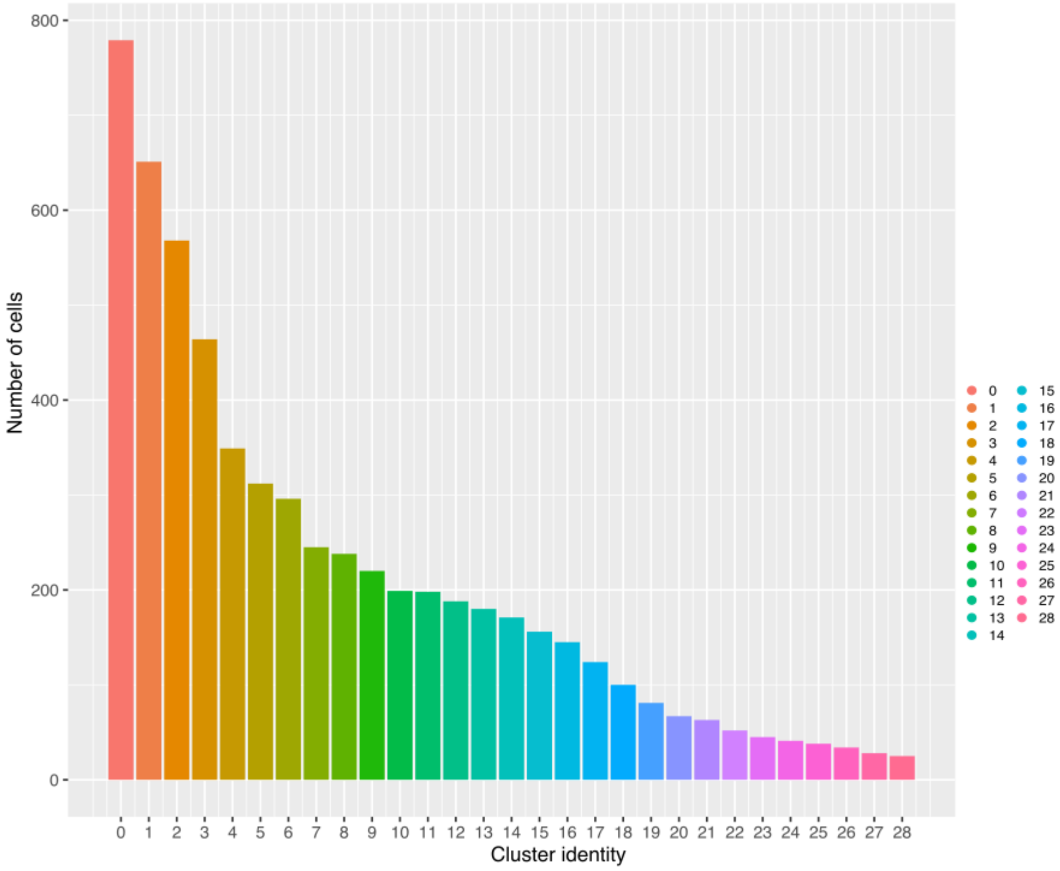


**Figure S3. The number of cells in each single-cell cluster of *B. diegensis.*** Cluster 0 had the highest cell count, and the lowest was Cluster 28. The color legend reflects the cluster identities.

**Cluster 14**

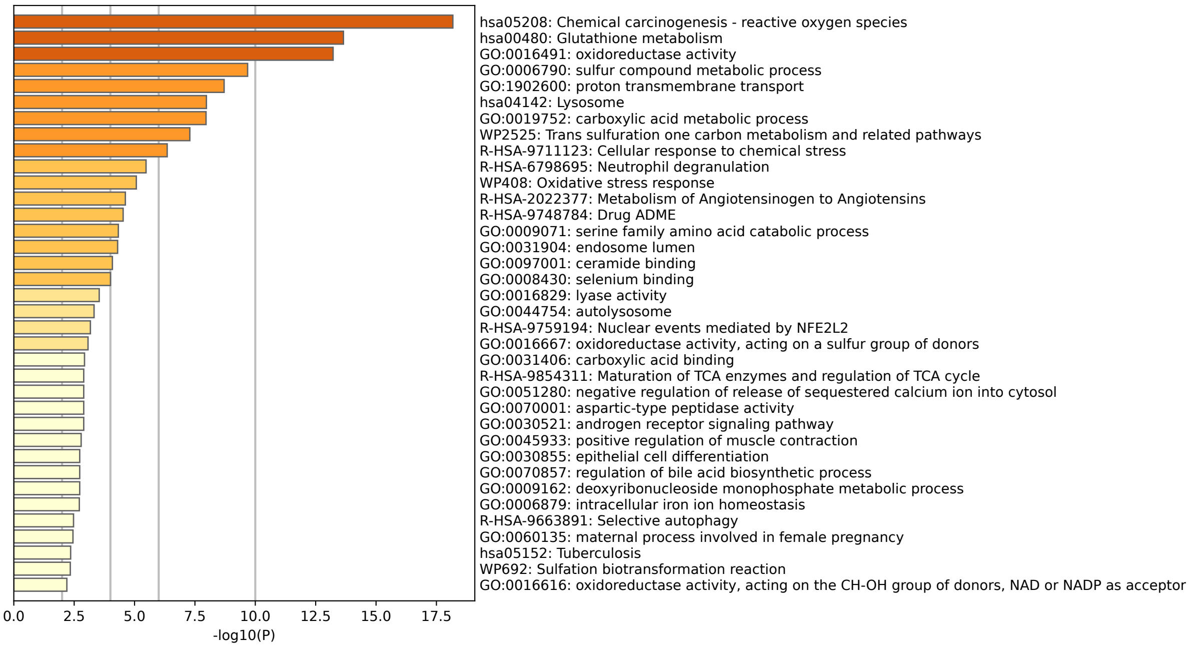


**Cluster 4**

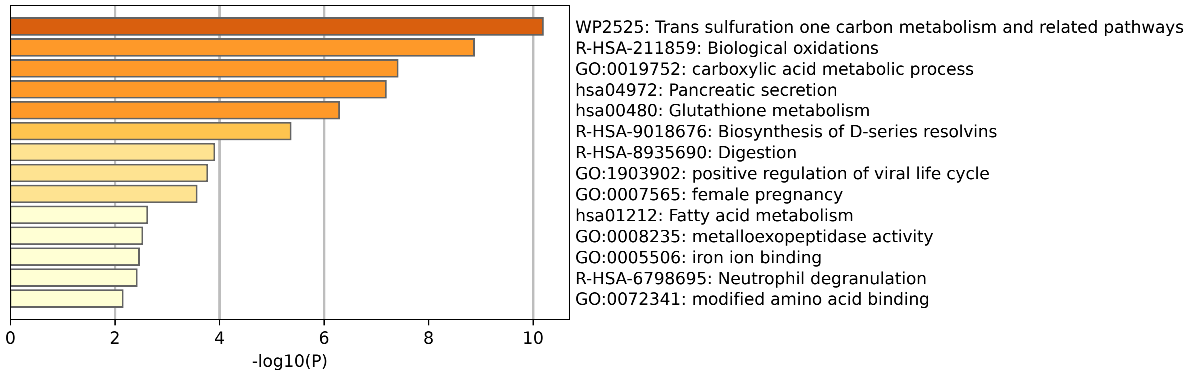


**Figure S4. Clusters 4 and 14.** The bar plots for Clusters 4 and 14 highlight the enriched biological processes and pathways associated with the top marker genes for these clusters. The x-axis represents -log10(P) values, indicating the statistical significance of each enrichment term. The longer the bar, the more significant is the enrichment.

**Cluster 10**

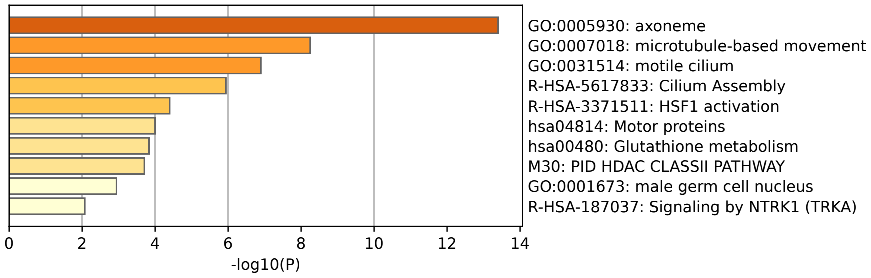


**Cluster 11**

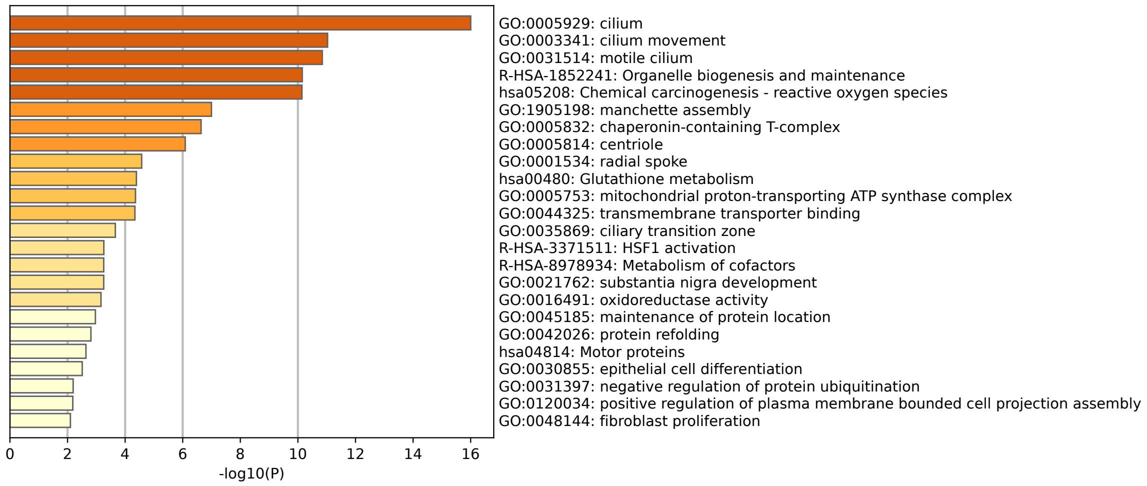


**Figure S5. Clusters 10 and 11.** The bar plots for Clusters 10 and 11 highlight the enriched biological processes and pathways associated with the top marker genes for these clusters. The x-axis represents -log10(P) values, indicating the statistical significance of each enrichment term. The longer the bar, the more significant is the enrichment.

**Cluster 0**

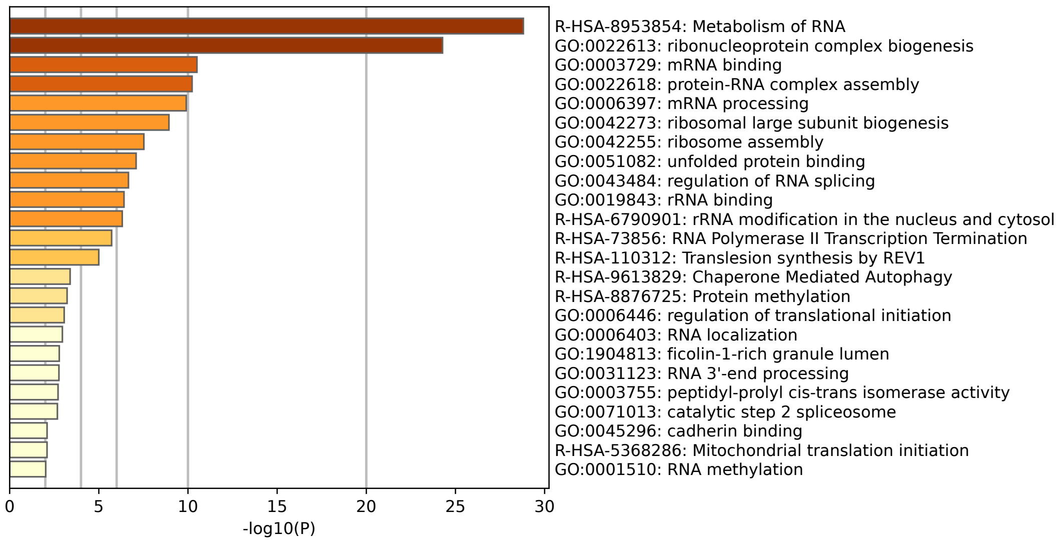


**Cluster 2**

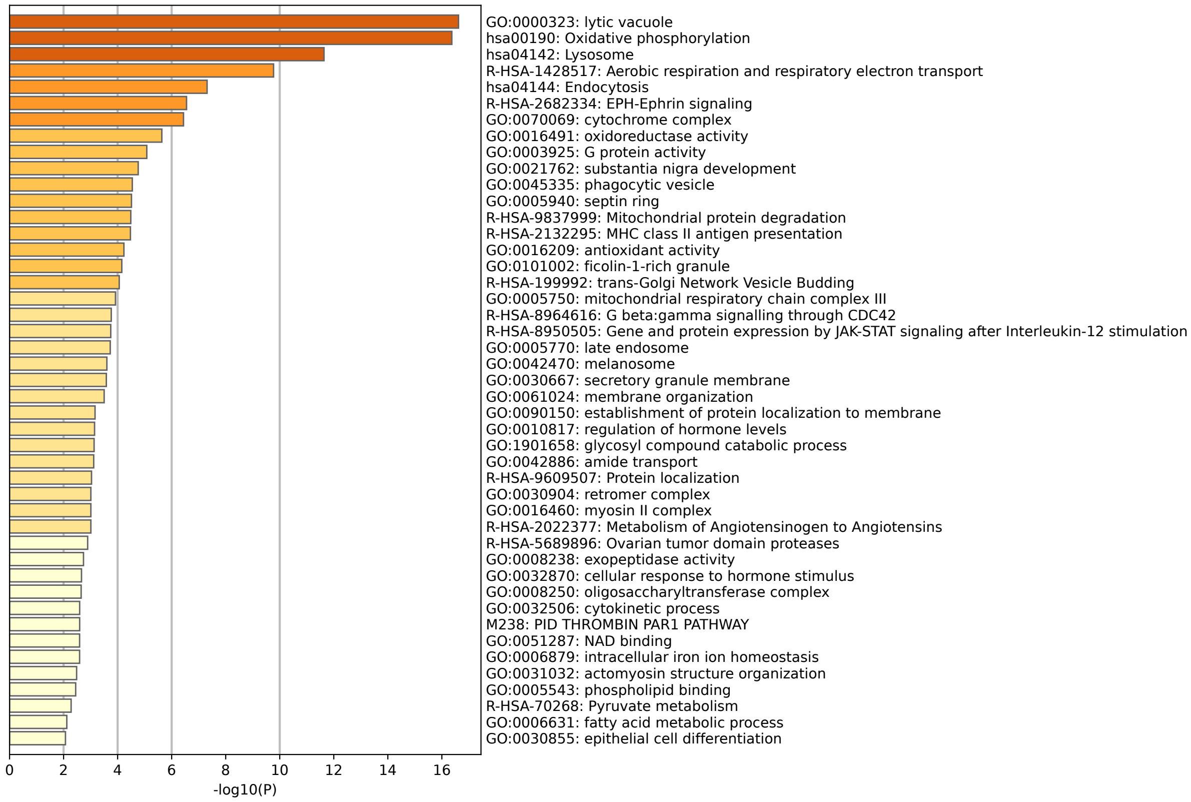


**Figure S6. Clusters 0 and 2.** The bar plots for Clusters 0 and 2 highlight the enriched biological processes and pathways associated with the top marker genes for these clusters. The x-axis represents -log10(P) values, indicating the statistical significance of each enrichment term. The longer the bar, the more significant is the enrichment.

Cluster 3

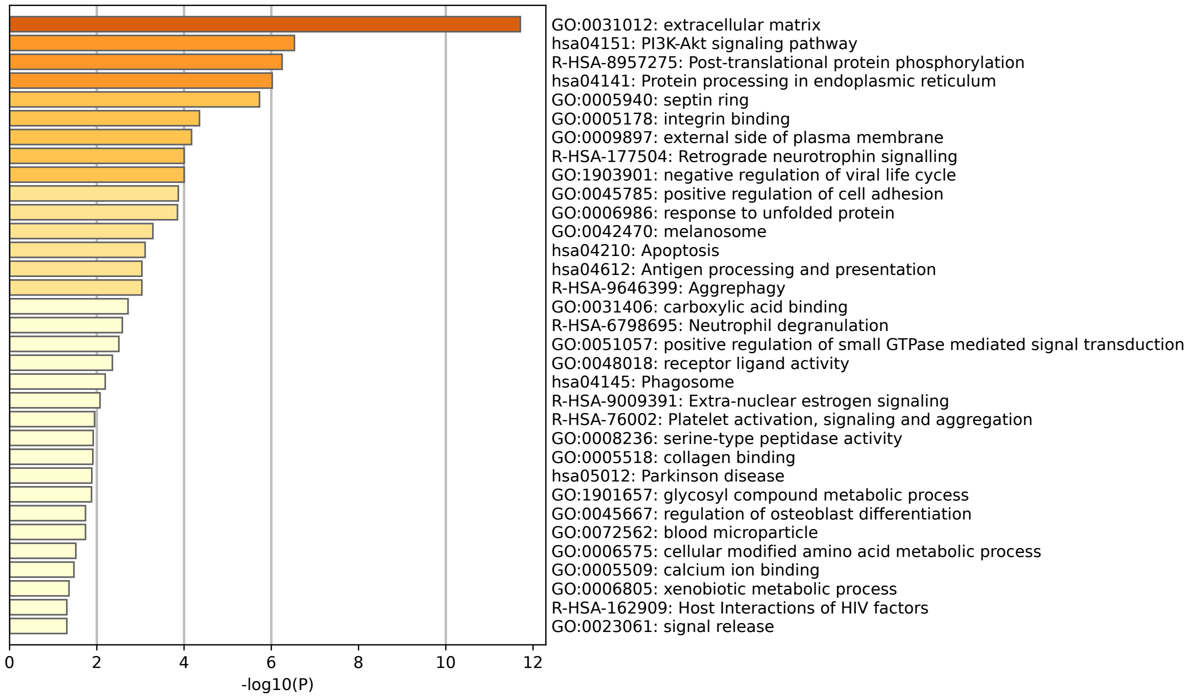


Cluster 6

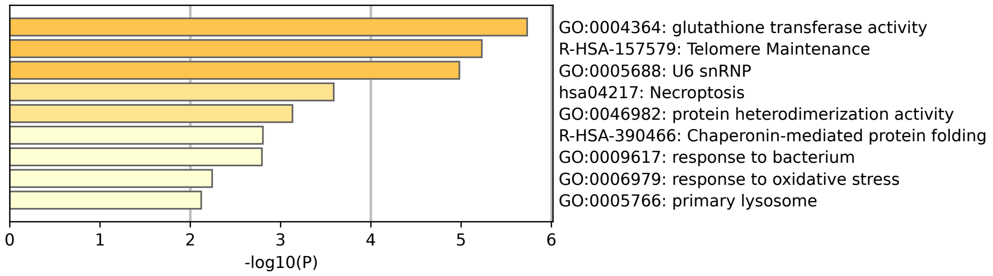


**Figure S7. Clusters 3 and 6.** The bar plots for Clusters 3 and 6 highlight the enriched biological processes and pathways associated with the top marker genes for these clusters. The x-axis represents -log10(P) values, indicating the statistical significance of each enrichment term. The longer the bar, the more significant is the enrichment.

### Cluster 9

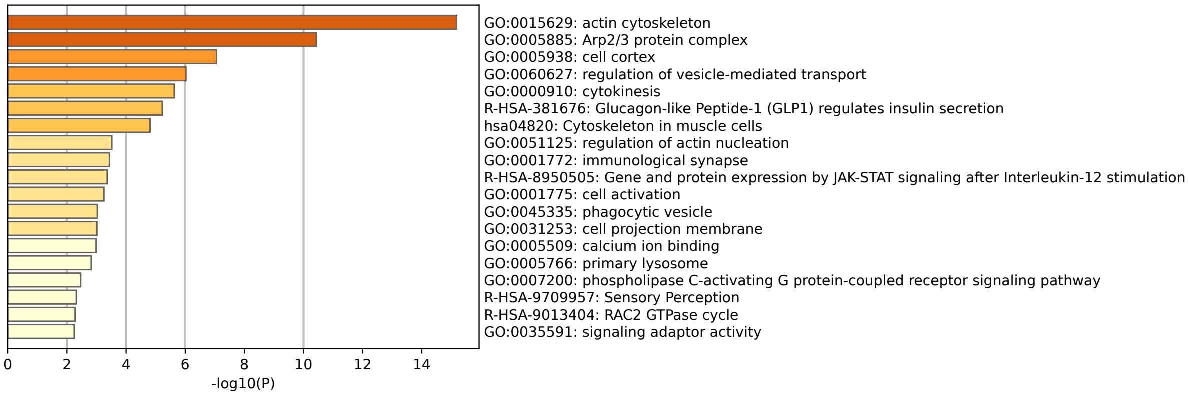


### Cluster 12

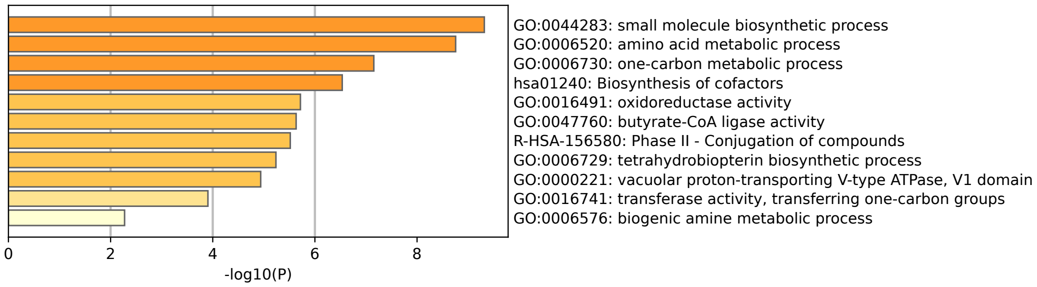


### Cluster 19

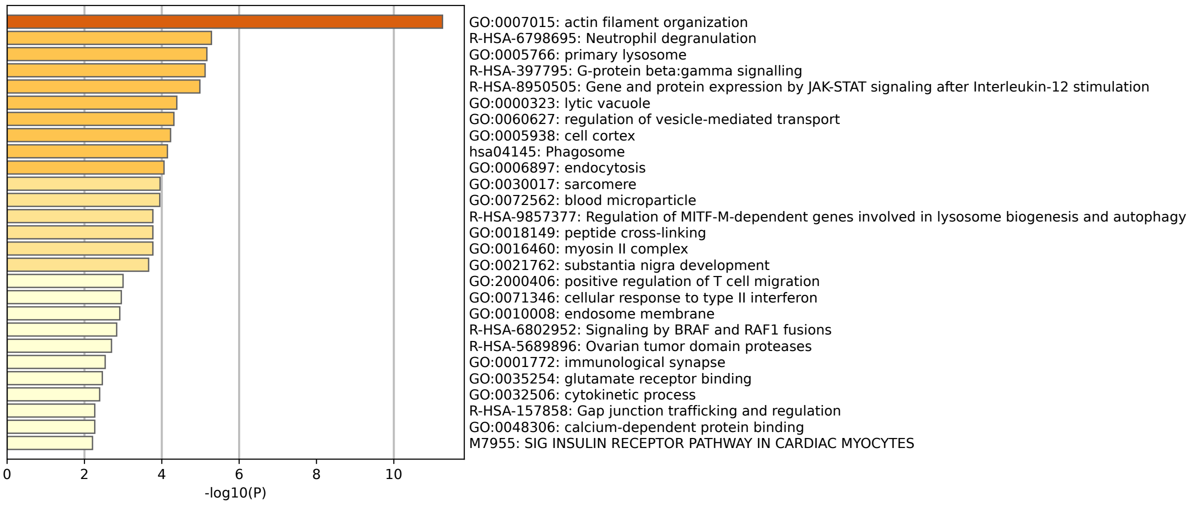


**Figure S8. Cluster 9, 19 and 12 cells.**

The bar plot for Clusters 9, 19, and 12 illustrates the enriched biological processes and pathways from the top marker genes for these clusters. The x-axis represents the -log10(P) values, reflecting the statistical significance of each enrichment term. The length of each bar indicates the significance level.

**Cluster 15**

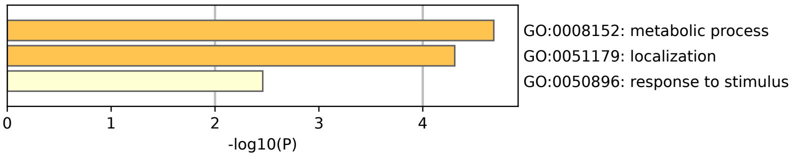


**Cluster 7**

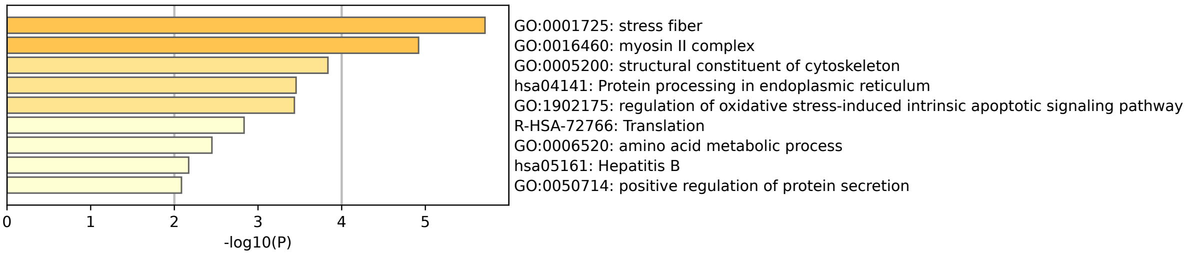


**Cluster 17**

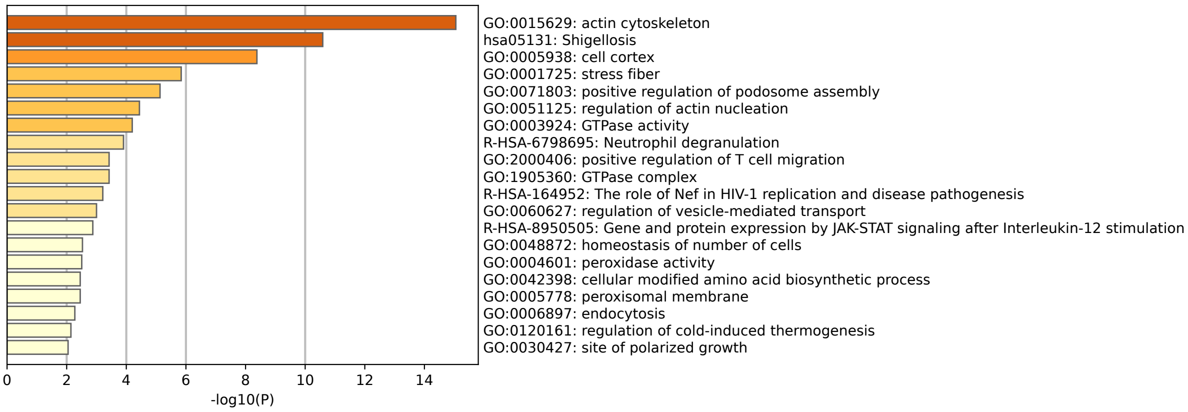


**Figure S9. Functional annotations of Clusters 15, 7 and 17.**

The bar plots for Clusters 15, 7, and 17 present the enriched biological processes and pathways identified from the top marker genes for these clusters. The x-axis shows the -log10(P) values, which measure the statistical significance of each enrichment term. The longer the bar, the more significant is the enrichment.

**Cluster 25**

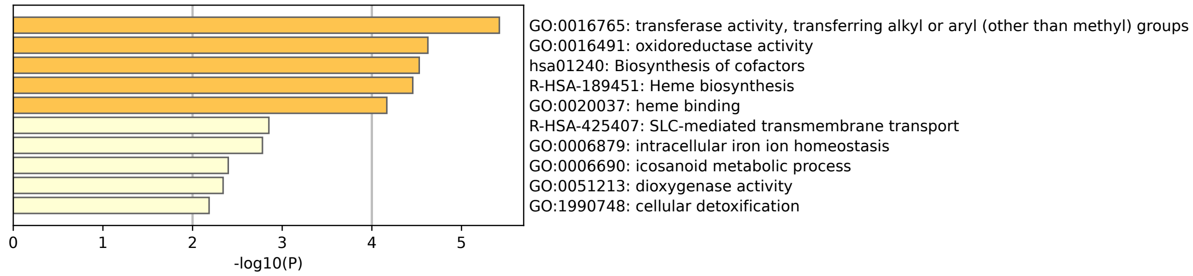


**Cluster 13**

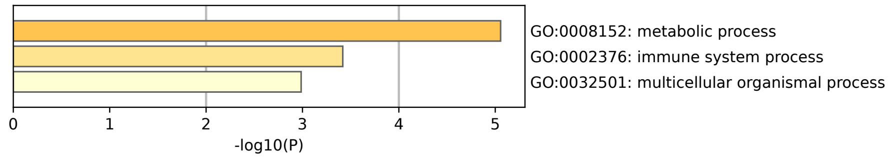


**Cluster 27**

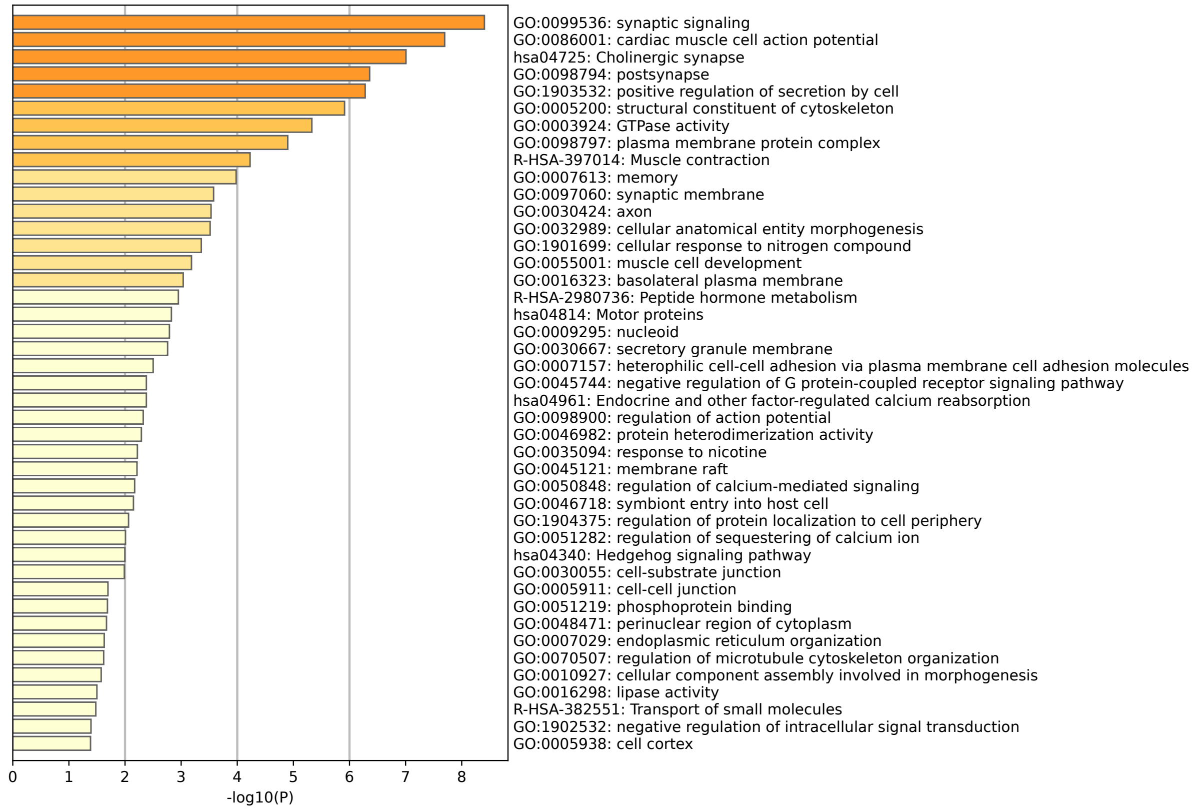


**Figure S10. Functional annotations of clusters 25, 27 and 13**

The bar plot for Clusters 25, 27 and 13 highlights the enriched biological processes and pathways associated with the top marker genes for these clusters. The x-axis represents -log10(P) values, indicating the statistical significance of each enrichment term.

.

\

**Cluster 8**

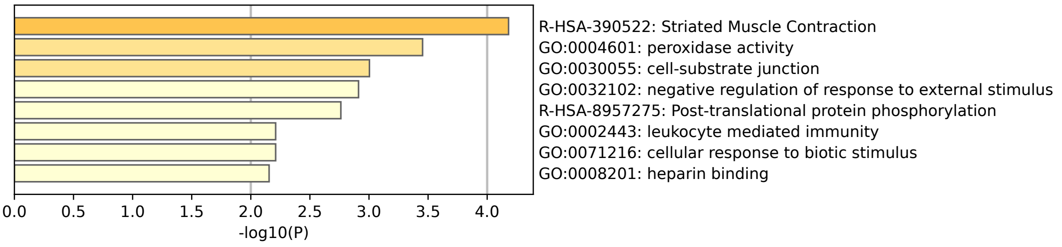


**Cluster 24**

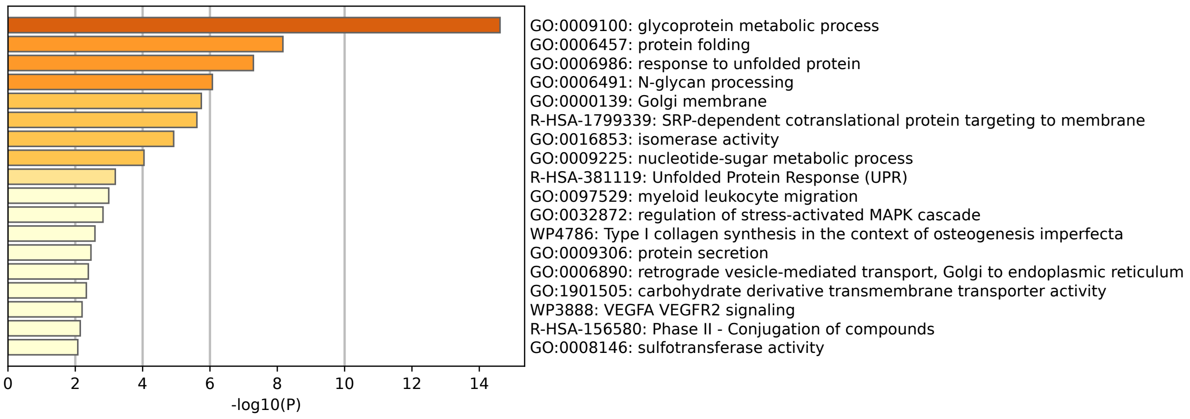


**Figure S11. Functional annotations of Clusters 8 and 24.** The bar plot for Clusters 8 and 24 highlights the enriched biological processes and pathways associated with the top marker genes for these clusters. The x-axis represents -log10(P) values, indicating the statistical significance of each enrichment term.

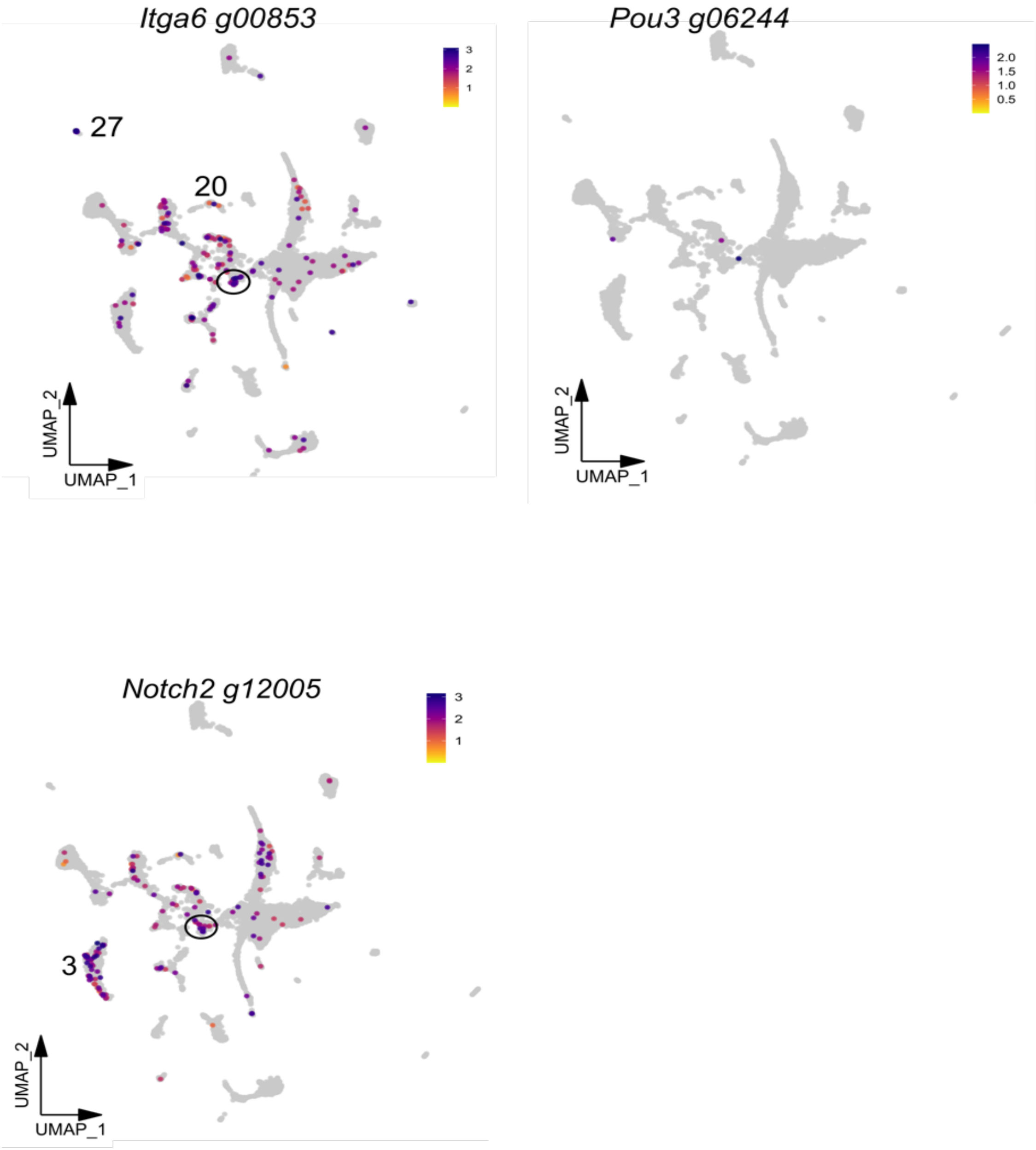


**Figure S12. Feature plots of genes previously linked to stemness in *Botrylloides* sp**. Circled cells are located within Cluster 6, are of interest.

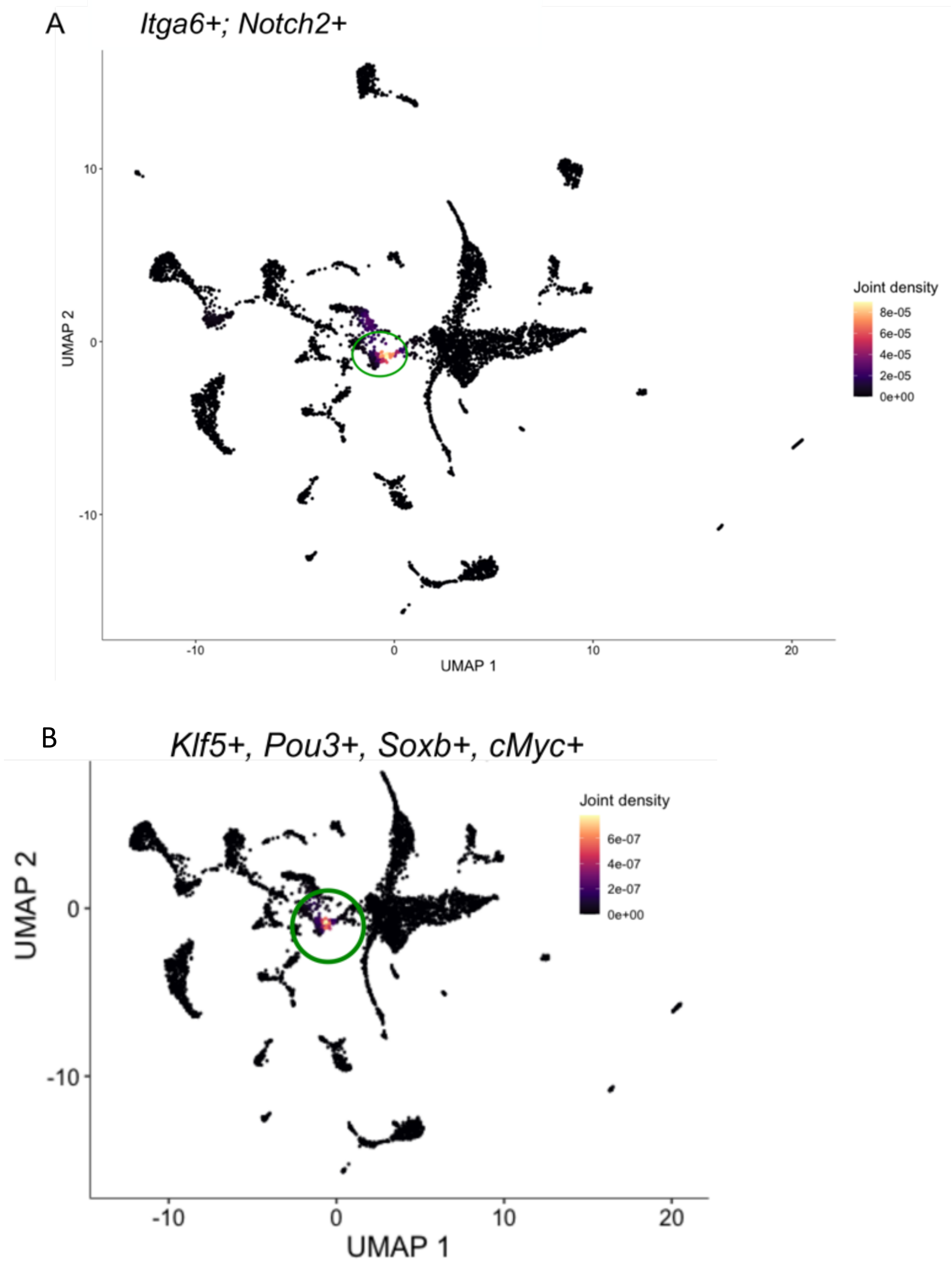


**Figure S13. Feature joint density plots. A**. Genes linked to stem cell function. **B**. Yamanaka factors. Circled cells are located within Cluster 6, which are of interest.

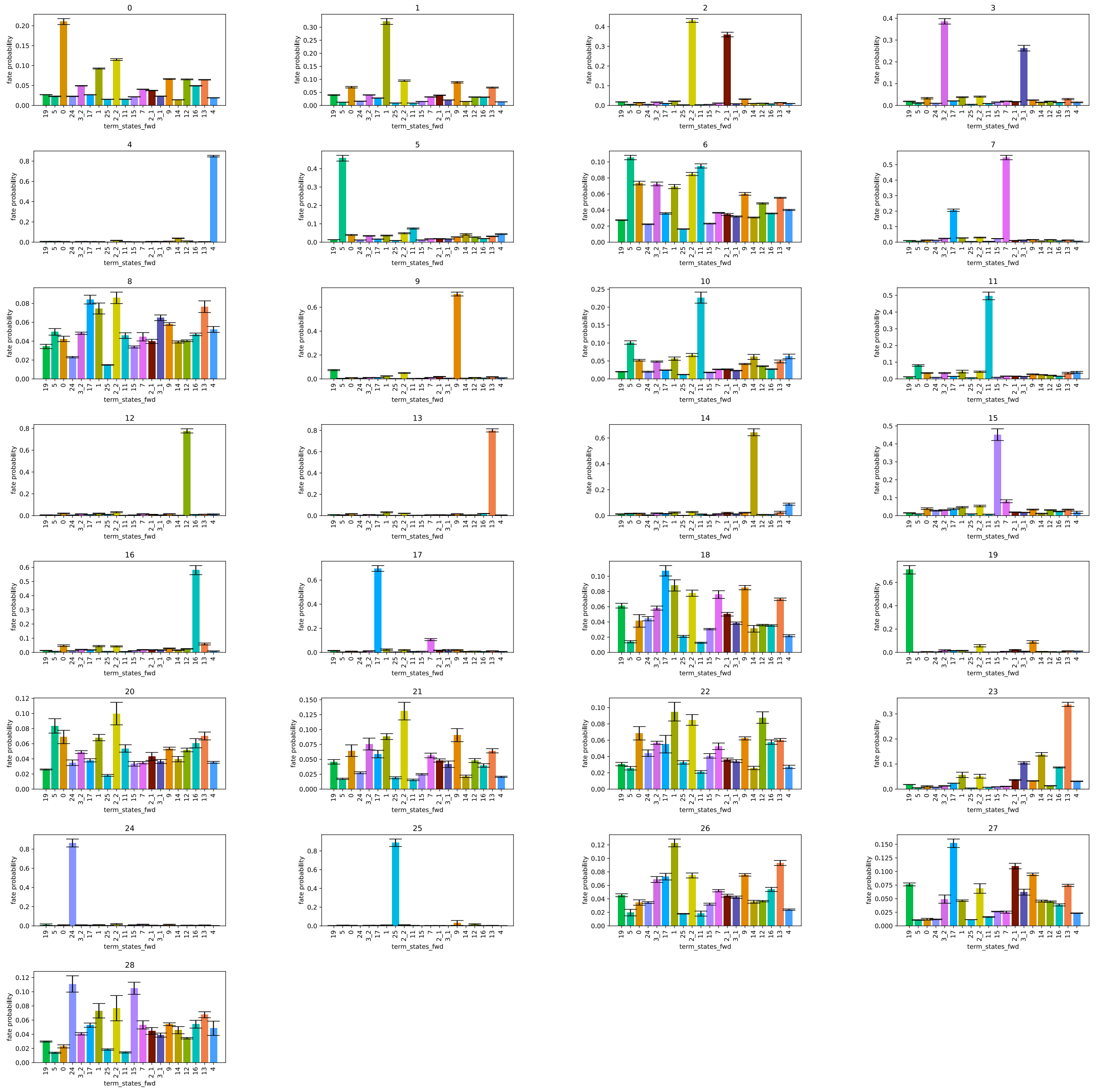


**Figure S14. CellRank cluster terminal fate probabilities**. The terminal fate probabilities for each cluster are shown, as determined by CellRank analysis. Each bar plot represents the probability distribution of the terminal states of the cells within a specific cluster. The x-axis of each bar plot denotes the various terminal fates, while the y-axis represents the probability of cells within each cluster obtaining each fate.

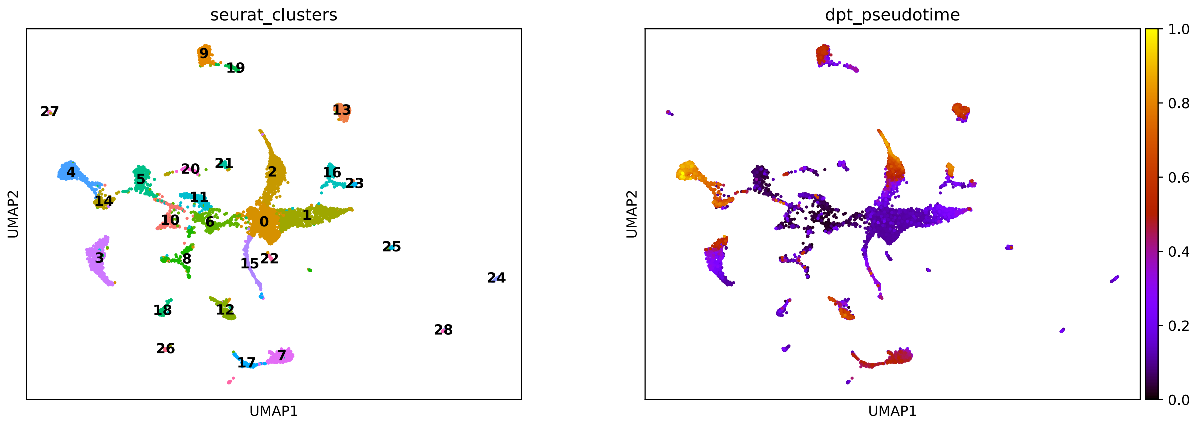


**Figure S15. UMAPs for diffusion pseudotime (dpt) and Seurat clusters.** Plots generated using CellRank (v. 2.0). The higher values (dpt_pseudotime) are later in the differentiation trajectory.
